# Adipose mitochondrial metabolism controls body growth by modulating cytokine and insulin signaling

**DOI:** 10.1101/2021.04.12.439566

**Authors:** Shrivani Sriskanthadevan-Pirahas, Michael J Turingan, Joel S Chahal, Erin Thorson, Savraj S Grewal

## Abstract

Animals need to adapt their growth to fluctuations in nutrient availability to ensure proper development and survival. These adaptations often rely on specific nutrient-sensing tissues and their control of whole-body physiology through inter-organ communication. While the signaling mechanisms that underlie this communication are well studied, the contributions of metabolic alterations in the nutrient-sensing tissues are less clear. Here, we show how reprogramming of adipose mitochondrial metabolism controls whole-body growth in *Drosophila* larvae. We find that dietary nutrients alter fat body mitochondrial morphology to lower their bioenergetic activity, which we see can rewire fat body glucose metabolism. Strikingly, we find that genetic reduction of mitochondrial bioenergetics just in the fat body is sufficient to accelerate body growth and development. These growth effects are caused by inhibition of the fat-derived adipokine, TNFα/Eiger, which leads to enhanced systemic insulin signaling, the main hormonal stimulator of body growth. Our work reveals how reprogramming of mitochondrial metabolism in one nutrient-sensing tissue is able to couple whole body growth to nutrient availability.

## Introduction

Animals often grow and live in conditions where food abundance varies. Thus, as they develop through their embryonic and juvenile stages, they must coordinate their metabolism with fluctuations in nutrition in order to properly regulate their growth (Mirth et al., 2021). Defects in this coordination can impair development and can lead to growth disorders and lethality.

The nutritional control of whole-body growth relies in large part on networks of inter-organ communication (Boulan et al., 2015; Droujinine and Perrimon, 2016; Koyama et al., 2020). Often a particular tissue functions as a primary nutrient sensor that then signals to other tissues and organs to control systemic physiology. Thus, while all animal cells are individually capable of sensing nutrients, these organ-to-organ communication networks provide a way to achieve organism-wide coordination of metabolism and growth in response to changes in nutrition (Gillette et al., 2021).

*Drosophila* larvae have been an excellent model system to understand how inter-organ communication governs body growth (Texada et al., 2020). Over a 4-5 day period, larvae increase in mass over 200-fold before developing into pupae. This growth relies on dietary nutrients and is largely controlled by a conserved endocrine insulin/insulin-like growth factor pathway. In particular, several *Drosophila* insulin-like peptides (dILPs) are secreted from brain insulin producing cells (IPCs) into the hemolymph where they can activate a conserved PI3K-Akt pathway in all tissues to drive growth (Grewal, 2009, 2012). The fat body has emerged as a central tissue involved in coupling dietary nutrients to systemic insulin signaling (Andersen et al., 2013). The fat body is similar to mammalian liver and adipose tissue in that it stores and mobilizes glycogen and fat stores. But the fat body can also sense changes in dietary nutrients and then signal to the brain through different cytokines and secreted peptides, termed adipokines. In rich nutrient conditions, the fat body secretes adipokines (such as upd2, GBP 1 and 2, stunted and CCHamide 2) that promote dILP release, while suppressing the release of adipokines (such as the TNF-α homolog, Eiger) that function to inhibit dILP release (Agrawal et al., 2016; Delanoue et al., 2016; Koyama and Mirth, 2016; Rajan and Perrimon, 2012; Sano et al., 2015). However, upon nutrient deprivation or starvation, this profile of adipokine signaling is reversed leading to reduced dILP release, and as a result, decreased body growth. Each adipokine can respond to distinct dietary nutrients, often through conserved nutrient responsive signaling molecules such as glucagon, AMPK and the target-of-rapamycin (TOR) kinase (Agrawal et al., 2016; Ingaramo et al., 2020; Rajan et al., 2017). However, it is not clear if and how changes in fat body metabolism couple nutrients to the regulation of adipokines.

Mitochondria are the central metabolic regulators of all animal cells. The classic textbook role for mitochondria is as bioenergetic organelles that efficiently generate ATP through oxidative phosphorylation (OxPhos) and the electron transport chain (ETC). However, mitochondrial metabolism also plays an important biosynthetic role by providing TCA cycle products as precursors to fuel cell metabolic processes such as generation of amino acids, lipids, and nucleotides.

In recent years, there has been surge of interest in exploring how these bioenergetic and biosynthetic functions are coupled to cell growth and proliferation. This interest has, in large part, been driven by the finding that many oncogenic signaling pathways can alter mitochondrial function in order to remodel cellular metabolism. These effects are thought to underline the concept of ‘Warburg cancer metabolism’, in which cancer cells can switch to glycolytic metabolism to generate ATP and use both glycolysis and the TCA cycle to generate biosynthetic precursors to promote their growth and proliferation (DeBerardinis and Chandel, 2016, 2020). These metabolic changes are also seen in proliferating cells during development (Miyazawa and Aulehla, 2018). It is also becoming clear that changes in mitochondrial metabolism do not just occur passively in response to alterations in cell growth, or that they function permissively to support the bioenergetic requirements of proliferating cells. Rather, mitochondrial metabolism can actively drive changes in cellular behaviors such as growth, proliferation and differentiation. For example, changes in mitochondrial activity can control the switch between proliferation and differentiation in stem cells (Homem et al., 2015; Khacho et al., 2016; Schell et al., 2017; Senos Demarco et al., 2019). Also, the inflammatory and secretory activity of immune and adipose cells can be directly controlled by changes in their mitochondrial metabolism (Mills et al., 2016; Ryan et al., 2019; Ryan and O’Neill, 2020; Tannahill et al., 2013).

While much of what we are learning about how mitochondrial metabolism drives growth comes from excellent studies in cultured cells, the mechanisms that operate *in vivo* in developing animals remain to be determined. Here, proper body growth is not simply a matter of increased cellular proliferation, but also requires coordination of tissue growth across multiple organs to ensure proportional size control. In this paper, we have used *Drosophila* larvae to explore how mitochondrial metabolism regulates animal growth and development. We have discovered a mechanism through which alterations in fat body mitochondrial bioenergetics can control whole-body growth via fat-body adipokines and systemic insulin signaling.

## Results

### Fat body mitochondrial architecture changes over the larval growth phase

Mitochondria consist of an outer membrane and inner membrane. The inner membranes form invaginations into the matrix known as cristae, which are the sites of the electron transport chain (ETC) complexes. Efficient mitochondrial OxPhos and ATP synthesis rely on the exquisite membrane organization of these mitochondrial cristae (Hackenbrock, 1968; John et al., 2005). To investigate the role of mitochondria in the fat body during larval growth phase, we first examined mitochondrial architecture in fat body cells at three different stages of larval development – 72hr and 96hr after egg-laying (AEL), which correspond to the growth phase of the larval period, and 120hr AEL, which corresponds to the end of the larval period when growth is slowed before metamorphosis into the pupal stage. We examined mitochondria by staining with MitoTracker Red staining and visualizing by confocal microscopy, and by transmission electron microscopy (TEM). Using both methods, we found fat body mitochondrial morphology changed dramatically over larval development (**Figures 1A-1C**). At 72 hrs. and 96 hrs. AEL, we observed large mitochondria with few cristae, suggesting low bioenergetic activity. However, at 120 hrs. AEL mitochondria were smaller with a denser, tight cristae structure as seen by the highly electrondense mitochondria in TEM images and by a marked increase in the intensity of the MitoTracker Red staining in the confocal image of the fat cells (**Figure 1C**). These changes are consistent with an increase in bioenergetic activity. These results indicate that fat body mitochondria change their size and morphology over the course of larval development, with features of lower bioenergetic activity predominating through the rapid growth phase and higher bioenergetic activity at the end of the growth period.

**Figure 1.**
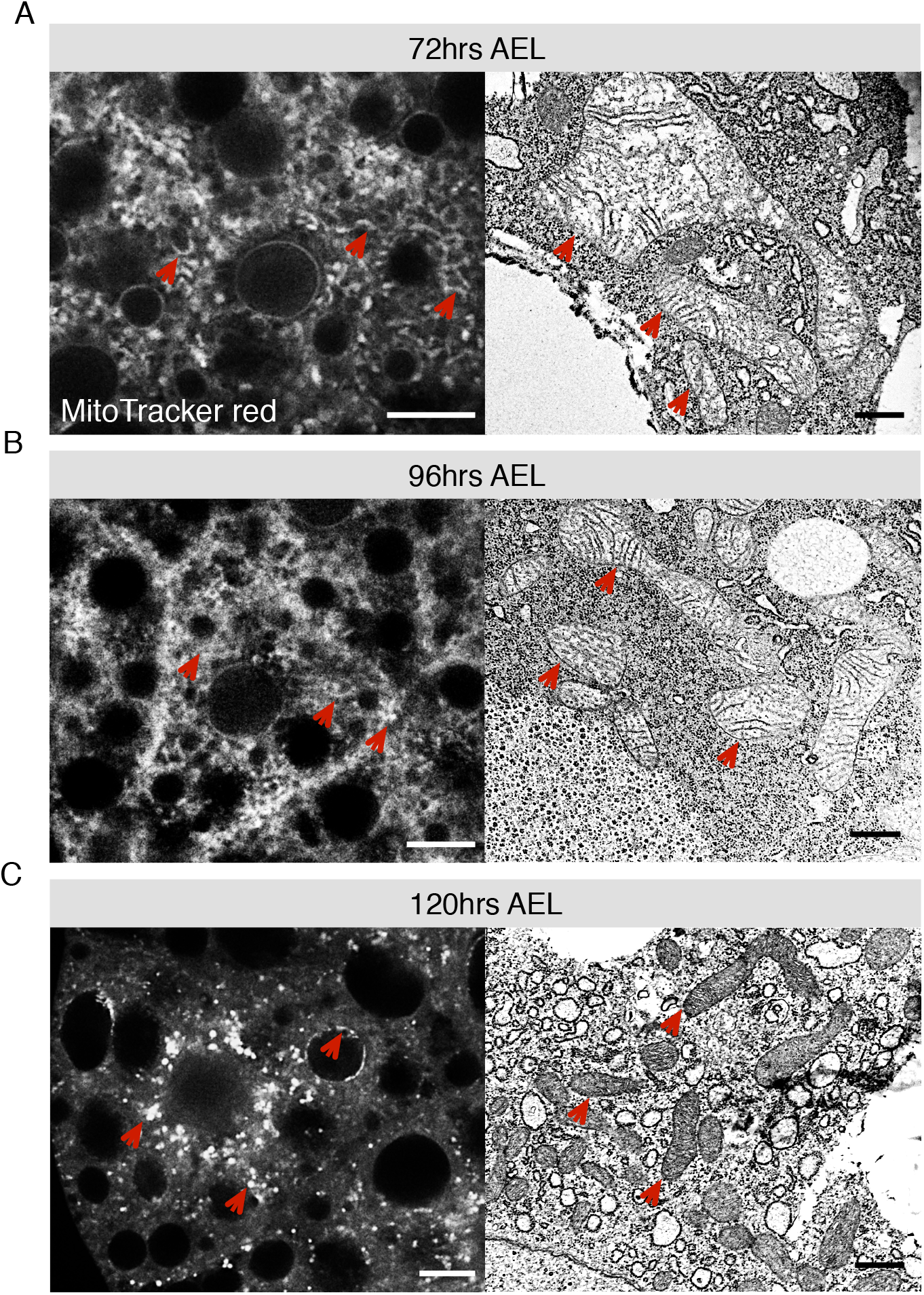
Fat body mitochondrial architecture changes over larval growth phase. (**A-C**) Representative confocal and electron micrographs of fat body mitochondria from larvae at 72 hrs AEL (A), 96 hrs AEL (B), and 120 hrs AEL (C) are shown. Confocal images of fat body were stained with MitoTracker Red. Red arrows indicate the mitochondria. The scale bars for confocal images represent 10 μm. TEM images of fat body mitochondria at 72 hrs AEL show larger mitochondria with sparse cristae formation. At 96 hrs AEL, fat body mitochondria are smaller compared to 72 hrs but still consist of low number of cristae. At the end of growth period (120 hrs AEL), fat body mitochondria become smaller in size with dense cristae formation. The scale bars for electron micrographs represent 500 nm.

### Nutrient availability regulates fat body mitochondrial activity

The rapid growth during the larval period is promoted in large part by dietary nutrients. To investigate whether nutrients influence mitochondrial function during the larval growth phase, we compared mitochondrial morphology in larvae growing in high vs. low nutrient conditions. To do this, we raised larvae on our normal diet until 72hrs. AEL and then switched them to vials containing either our normal lab food (normal food) or our lab food diluted to 20% with water and agar (low food) (**Figure S1A**). Under these low food conditions, both growth and development are slowed and, as a result larvae took longer to develop to pupae (**Figure 2A, Figure S1B**) and had a smaller final body size (**Figure 2B**), although overall survival was not strongly affected (**Figure S1C**). However, unlike complete nutrient deprivation, our low nutrient diet did not induce autophagy, indicating that we are not inducing starvation stress conditions (**Figure S2**).

**Figure 2.**
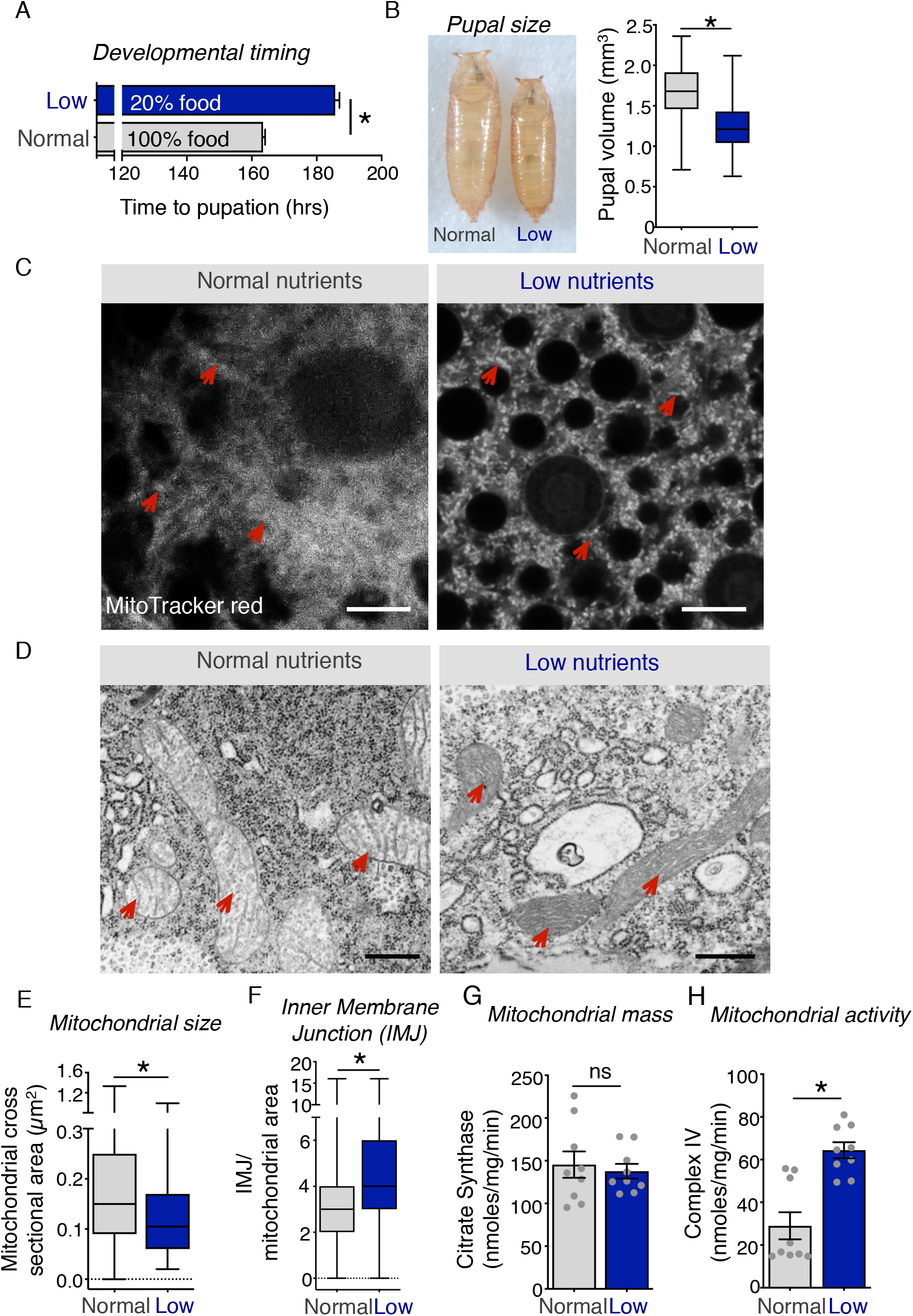
Nutrient availability regulates fat body mitochondrial activity. (**A**) Larvae grown on normal food were transferred to diluted food (20% of the normal food) and time to pupation was calculated and plotted as bar graph. Data are presented as mean +/− SEM (*p < 0.05, Mann-Whitney U test). (**B**) Pupal sizes were measured for the larvae grown on normal food (gray label) and 20% food (blue). Representative images are shown, and the quantitative data represents mean ± +/− SEM (*p < 0.05, unpaired t-test with welch correction). (**C**) Fat bodies were dissected from 96 hrs AEL larvae maintained in normal food or on low nutrient food from 72 hrs to 96 hrs. Fat bodies were then stained with MitoTracker Red and imaged with confocal microscopy. Representative images are shown. Arrows indicate the mitochondria. Scale bar represents 10 μm. (**D**) Electron micrographs show the fat body mitochondrial ultrastructure in normal food vs low nutrient conditions. Mitochondria from larvae grown on low nutrient food have denser cristae compared to the control. Red arrows indicate the mitochondria. Scale bar represents 500 nm. (**E**) Mitochondrial cristae density is quantified using electron micrographs images in low nutrient fat body vs control. NIH image J software was used to quantify the inner membrane junctions per mitochondrial cross sectional area. Data are presented as box plots (25%, median and 75% values) with error bars indicating the min and max values (*p < 0.0001, unpaired t-test, n > 200 mitochondria). (**F**) Fat body mitochondrial cross sectional area is higher in larvae grown in normal food compare to the larvae from low nutrient condition. Mitochondrial cross sectional area was calculated using NIH ImageJ software based on electron micrographs. Data are presented as box plots (25%, median and 75% values) with error bars indicating the min and max values (*p < 0.0001, unpaired t-test, n > 250 mitochondrial cross sectional area). See also Figure S1 and S2. (**G**) Mitochondrial citrate synthase activity in larvae grown on normal food vs low nutrient food. Mitochondria were isolated from larvae at 96 hrs AEL from normal vs low nutrient condition. Citrate synthase activity was measured using a biochemical assay and used as a marker for mitochondrial mass. Data are represented as mean ± SEM, with individual data points plotted as symbols. (ns, not significant p>0.05, unpaired t-test) (**H**) Mitochondrial OxPhos complex IV activity is higher in larvae grown on low nutrient condition compared to control animals. Complex IV activity was measured in isolated mitochondria from whole larvae and normalized to total protein concentration and mitochondrial mass. Data are represented as mean ± SEM, with individual data points plotted as symbols (*p < 0.05, unpaired t-test). See also Figures S1-S3.

We then compared fat body mitochondria in larvae grown on normal vs. low food conditions. We observed that in normal food at 96 hrs. AEL (when larvae are growing rapidly), fat body mitochondria were fused with sparse cristae formation (**Figure 2C and 2D**). In contrast, when larvae were switched to low nutrient conditions from 72 - 96 hrs, mitochondria appeared smaller with tight, dense cristae (**Figure 2C and 2D**). We quantified TEM images and found that fat body mitochondrial size was lower in low nutrient food (**Figure 2E and S3B**). In addition, we quantified mitochondrial cristae density by measuring the inner membrane junctions (IMJ) per mitochondrial cross section images and found that there was an increase in IMJ in low nutrient conditions (**Figure 2F**). Moreover, when we binned mitochondria into three groups based on their cristae morphology, we saw that a higher percentage of mitochondria from low food conditions were in the dense cristae group, whereas the majority of mitochondria from fat bodies of larvae in normal food were in the sparse cristae group (**Figure S3A**).

Tight cristae are an indication of highly respiring mitochondria. Therefore, we investigated whether mitochondrial OxPhos activity was altered under low nutrient conditions as suggested by the changes in fat body mitochondrial ultrastructure. We found that in low nutrient conditions, isolated larval mitochondria had higher complex IV activity, while the mitochondrial mass was unaffected (**Figure 2G and 2H**). Taken together these results indicate that in high nutrient conditions, mitochondria have low bioenergetic activity and are likely engaged in more biosynthetic activity, but upon switching to low nutrient conditions, fat body mitochondria increase their cristae density and show higher bioenergetic activity.

### Lowering fat body mitochondrial bioenergetic activity accelerates growth and development

We next wanted to explore the significance of these changes in fat body mitochondrial metabolic activity. To do this we used RNAi to knock down the mitochondrial transcription factor A, TFAM. TFAM is a nuclear encoded factor that localizes to mitochondria to specifically transcribe the mitochondrial genome, including the 13 essential genes encoding for ETC complex proteins (Kang et al., 2007). We found that whole-body knockdown of TFAM (using the *da-GAL4* driver) led to reduced mitochondrial gene expression (**Figure S4**) as well as reduced mitochondrial complex IV activity, as previously seen with TFAM knockdown in mice (Hamanaka et al., 2016), without a significant change in mitochondrial mass **(Figure 3A, B)**.

**Figure 3.**
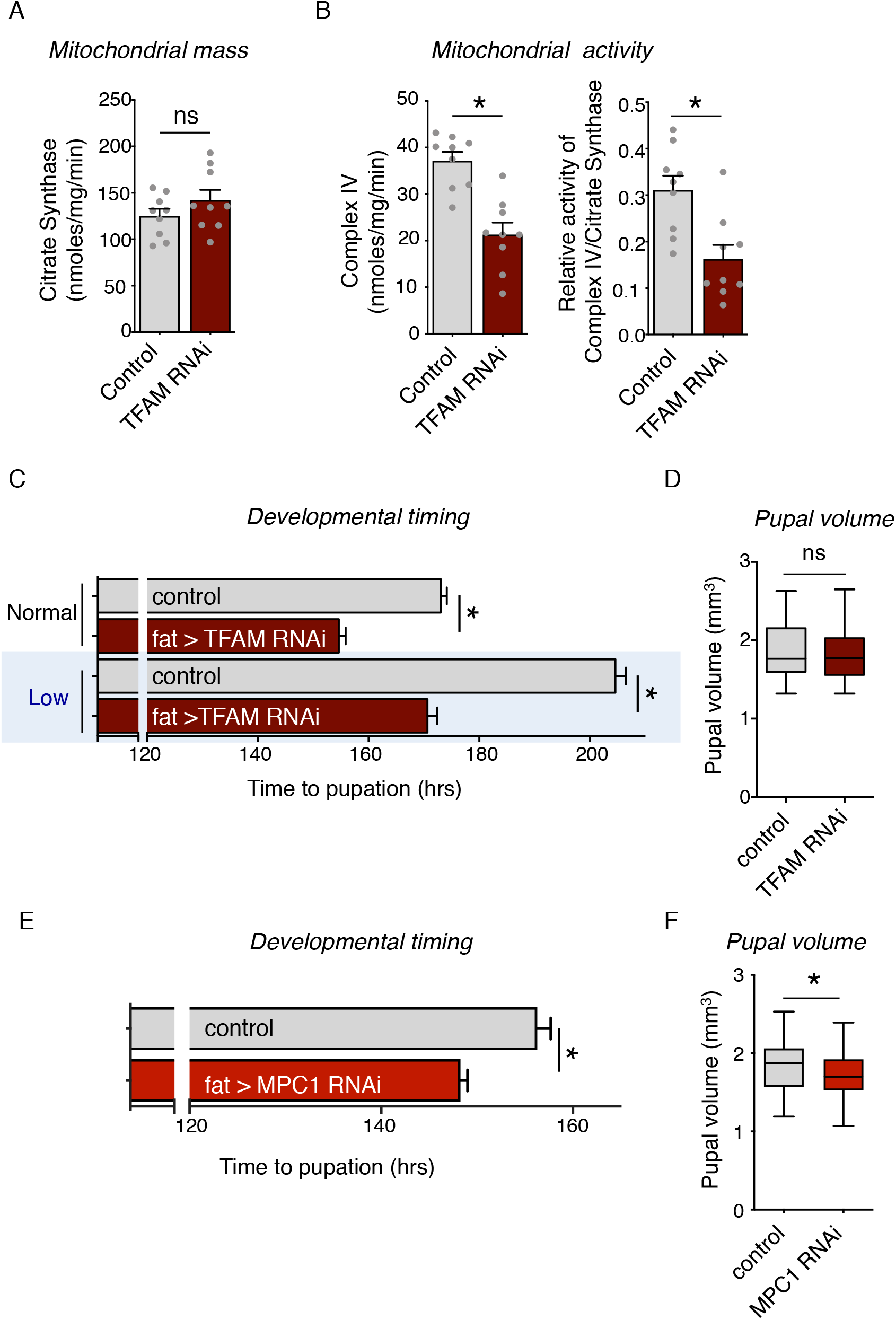
Lowering fat body bioenergetic activity accelerates growth and development. (**A**) TFAM knockdown has no effect on citrate synthase activity. Control larvae (*da > +*) or larvae expressing an inverted repeat RNAi transgene to TFAM (*da > TFAM RNAi*) were maintained in normal food from hatching to 96 hrs of development. Then, mitochondria were isolated from the larvae and biochemical assays were performed to measure the complex IV activity. Data are represented as mean ± SEM, with individual data points plotted as symbols (ns = not significant, p>0.05, unpaired t-test). (**B**) TFAM knockdown leads to reduced mitochondrial complex IV activity. Mitochondrial complex IV activity in control (*da > +*) vs TFAM RNAi (*da > TFAM RNAi*) larvae was measured and normalized to total protein concentration and mitochondrial mass. Data are represented as mean ± SEM, with individual data points plotted as symbols (*p < 0.05 unpaired t-test). (**C**) Fat-specific TFAM RNAi accelerates development both in normal and low nutrient condition. Time to pupation was measured in control (*r4 > +*) vs TFAM RNAi (*r4 > UAS-TFAM RNAi*) larvae. Data are represented as mean ± SEM (*p < 0.05 Mann-Whitney U test, n > 200 animals). (**D**) Fat body specific TFAM knock down has no effect on pupal size. Pupal volume was measured in control (*r4 > +*) vs TFAM RNAi (*r4 > UAS-TFAM RNAi*) larvae. Data are presented as box plots (25%, median and 75% values) with error bars indicating the min and max values. (**E**) MPC1 knockdown in fat accelerates larval growth and development. Data represent mean ± SEM (*p < 0.05 Mann-Whitney U test, n >143 animals). Time to pupation was measured in control (*r4 > +*) vs MPC1 RNAi (*r4 > UAS-MPC1 RNAi*) larvae. (**F**) Fat body specific MPC1 knockdown leads to a small reduction in body size. Pupal volume was measured in control (*r4 > +*) vs MPC1 RNAi (*r4 > UAS-MPC RNAi*) larvae. Data are presented as box plots (25%, median and 75% values) with error bars indicating the min and max values. (*p < 0.05 unpaired t-test, n > 330 animals per condition). See also Figures S4-S7.

To examine the effects of lower fat body OxPhos, we then use the fat body driver, *r4-GAL4,* to direct TFAM RNAi specifically to the fat body. Strikingly, we found that fat body TFAM knockdown led to a significant acceleration in development, with larvae taking a shorter time to reach the pupal stage (**Figure 3C**). Furthermore, this faster development was accompanied by an increase in growth rate since these larvae reached the same final pupal size as control animals (**Figure 3D**). We also found that fat body TFAM knock down could reverse the slower development of larvae growing in low nutrient conditions (**Figure 3C**). These effects were also observed with two additional independent TFAM RNAi lines (**Figure S5 and S6**), but were not seen in transgenic flies carrying a UAS-TFAM RNAi transgene alone (**Figure S7**). To explore these findings further, we used an alternate method to lower mitochondrial bioenergetic activity. We chose to target the mitochondrial pyruvate carrier 1 (MPC1) which is essential for transporting pyruvate into mitochondria for OxPhos (Bricker et al., 2012). We found that fat body specific MPC1 RNAi also led to accelerated larval growth and development, mimicking the effects of TFAM RNAi (**Figure 3E and 3F**). Taken together, our results demonstrate that lowering mitochondrial bioenergetic activity in the larval fat body alone can accelerate the rate of animal growth and development.

### Fat body TFAM knockdown alters glucose metabolism

We next wanted to investigate how lowering OxPhos activity alters cell autonomous fat body function to drive body growth. To do this, we performed mRNA Seq analysis to examine the transcriptome of fat bodies from control vs. TFAM knock down larvae (**Figure S8–11**). RNA Seq differential expression analysis revealed a significant change in 1758 transcripts, with 1213 transcripts showing reduced expression levels and 545 transcripts with elevated expression levels. KEGG pathway and Gene Ontology analyses revealed that most of the downregulated genes were enriched in genes responsible for mitochondrial gene expression and oxidative phosphorylation, while the upregulated genes were enriched for metabolic genes (**Figure S8–11**). Interestingly, among these upregulated genes were those encoding for enzymes involved in different aspects of glucose metabolism (**Figure 4A and Figure S10**), including glycogenolysis, gluconeogenesis/glycolysis and the pentose phosphate pathway. When we examined fat body glycogen levels using Periodic Acid Schiff staining, we saw that TFAM knockdown fat bodies had lower glycogen levels (**Figure 4C and 4D**). We also found that glucose uptake was significantly reduced in fat body with TFAM knock down (**Figure 4E and 4F**). In contrast, we saw little effect on lipid droplet size (**Figure S12).** These findings suggest that TFAM knockdown leads to reprogramming of fat body glucose metabolism.

**Figure 4.**
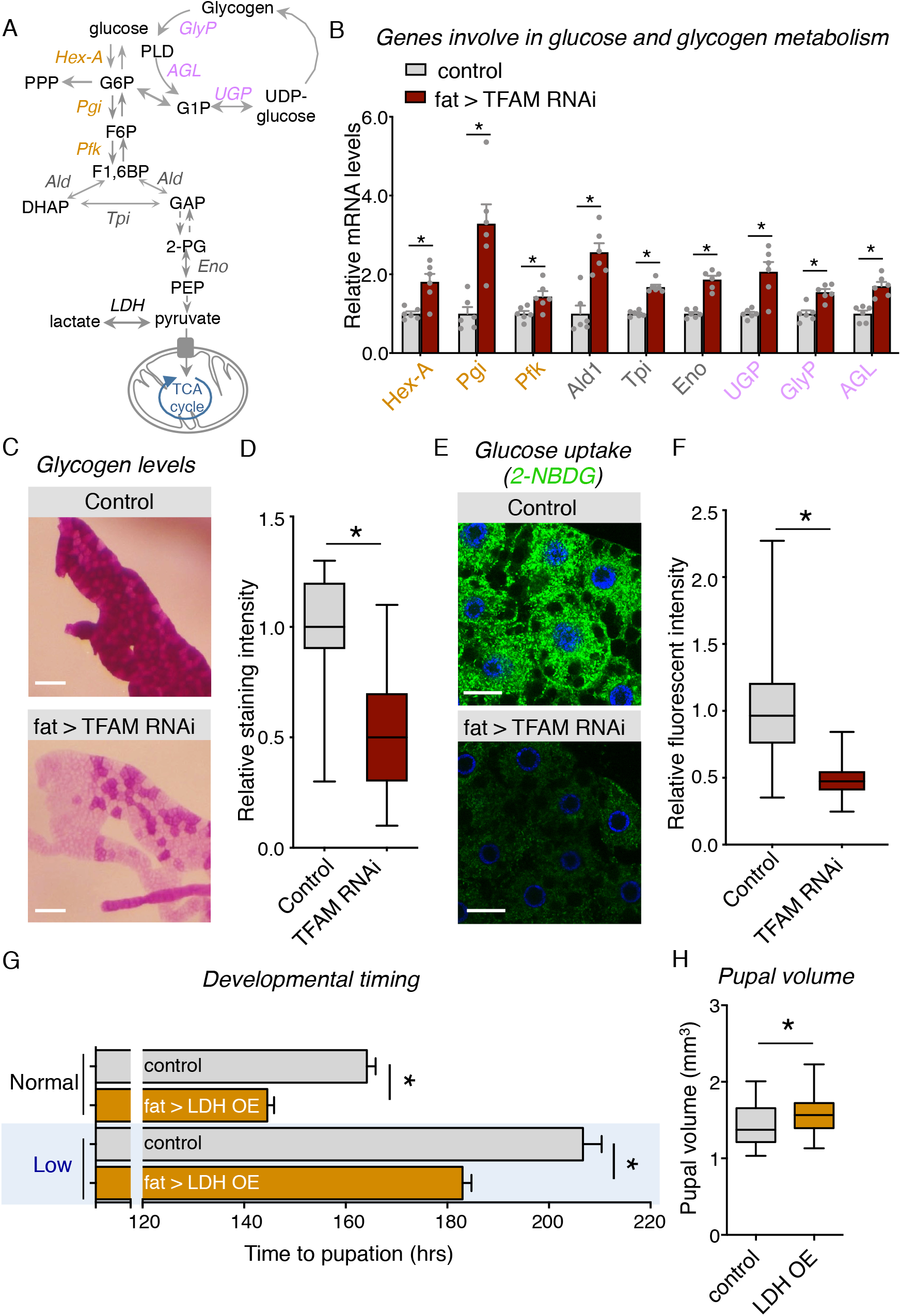
Fat body TFAM knock down alters glucose metabolism. (**A**) A schematic showing glucose and glycogen metabolic pathway. (**B**) Fat body mRNA-Seq analysis shows increase in genes involved in glucose and glycogen metabolism following TFAM knockdown. Fat bodies were dissected from control (*r4 > +*) vs TFAM RNAi (*r4 > UAS-TFAM RNAi*) larvae at 96 hrs AEL and total RNA was isolated for mRNA-Seq analysis. Data are represented as mean relative levels ± SEM with individual data points plotted as symbols (*p < 0.05 and ns = not significant, unpaired t-test, n = 6 per condition). (**C**) Dissected fat bodies were stained with Periodic Acid Schiff (PAS) staining to measure glycogen levels specifically in fat cells in control (*r4 > +*) vs TFAM RNAi (*r4 > UAS-TFAM RNAi*) larvae. Scale bar represents 500 μm. (**D**) Quantification of glycogen staining in (C). Data are presented as box plots (25%, median and 75% values) with error bars indicating the min and max values. (*p < 0.05 unpaired t-test). (**E**) Glucose uptake was analyzed with fluorescently labeled deoxy glucose, 2-NBDG. Fat bodies were dissected from control (*r4 > +*) vs TFAM RNAi (*r4 > UAS-TFAM RNAi*) larvae at 96 hrs AEL and stained with either 2-NBDG to measure glucose uptake. Scale bar represents 5 μm. (**F**) Quantification of glucose uptake in (E). Data are presented as box plots (25%, median and 75% values) with error bars indicating the min and max values (*p < 0.05 unpaired t-test). (**G**) Fat body overexpression of LDH accelerates animal growth and development in normal and low nutrient condition. Data represent mean ± SEM (*p < 0.05 Mann-Whitney U test, n > 80 animals per condition). (**H**) Fat body specific LDH overexpression leads to a slight increase in pupal size compared to controls. Time to pupation was measured in control (*r4 > +*) vs LDH overexpressing (OE) (*r4 > UAS-LDH*) larvae. Data are presented as box plots (25%, median and 75% values) with error bars indicating the min and max values (*p < 0.05 unpaired t-test, n > 200 animals).

To further explore the consequence of altered fat body glucose metabolism, we overexpressed lactate dehydrogenase (LDH). LDH is a key glucose metabolic enzyme that functions to catalyze the reversible conversion of lactate to pyruvate with the reduction of NAD+ to NADH and vice versa. Interestingly we saw that animals with fat LDH overexpression accelerated their growth and development both in normal and low nutrient conditions (**Figure 4G, 4H and S13**). Taken together, our results suggest that TFAM knockdown reprograms fat body glucose metabolism, and that these effects may contribute to stimulation of body growth.

### Fat body TFAM knockdown increases systemic insulin signaling via modulation of Eiger/TNFα signaling.

Our results indicate that lowered fat body mitochondrial bioenergetics can enhance whole-body growth and development. One established mechanism through which the fat body impacts body growth is through the endocrine control of insulin signaling, the main regulator of growth in larvae (Texada et al., 2020). In particular, fat-to-brain signaling promotes the release of three dILPs (2, 3 and 5) from the IPCs into the larval hemolymph. Here, the dILPs can bind to cell surface insulin receptors on all peripheral tissues to stimulate a conserved PI3K-Akt pathway that inhibits the nuclear localization and transcriptional activity of the fork head transcription factor, FOXO. We therefore examined whether fat body TFAM knockdown alters whole-body insulin signaling. We found that fat specific TFAM RNAi led to reduced whole-body expression levels of three FOXO target genes, 4E-BP, dILP6 and InR, compared to control larvae. (**Figure 5A**). In addition, we found that whole-body levels of phosphorylated AKT were increased following fat body-specific TFAM knockdown (**Figure 5B and 5C**). These results indicate that systemic insulin signaling is increased following reduction in fat body mitochondrial OxPhos activity. To explore, whether these effects were involved in the stimulation of growth seen upon fat body TFAM knockdown, we examined the effects of lowering insulin signaling. To do this, we used a genetic deficiency that removes the three IPC-expressed dILPs - 2, 3 and 5. We found that, in contrast to the growth stimulatory effects seen in a wild-type background **(Figure 3C),** when we knockdown TFAM in the fat bodies of larvae heterozygous for the dILP deficiency, there was no increase in either development rate **(Figure 5D)** or body growth **(Figure 5E).**

**Figure 5.**
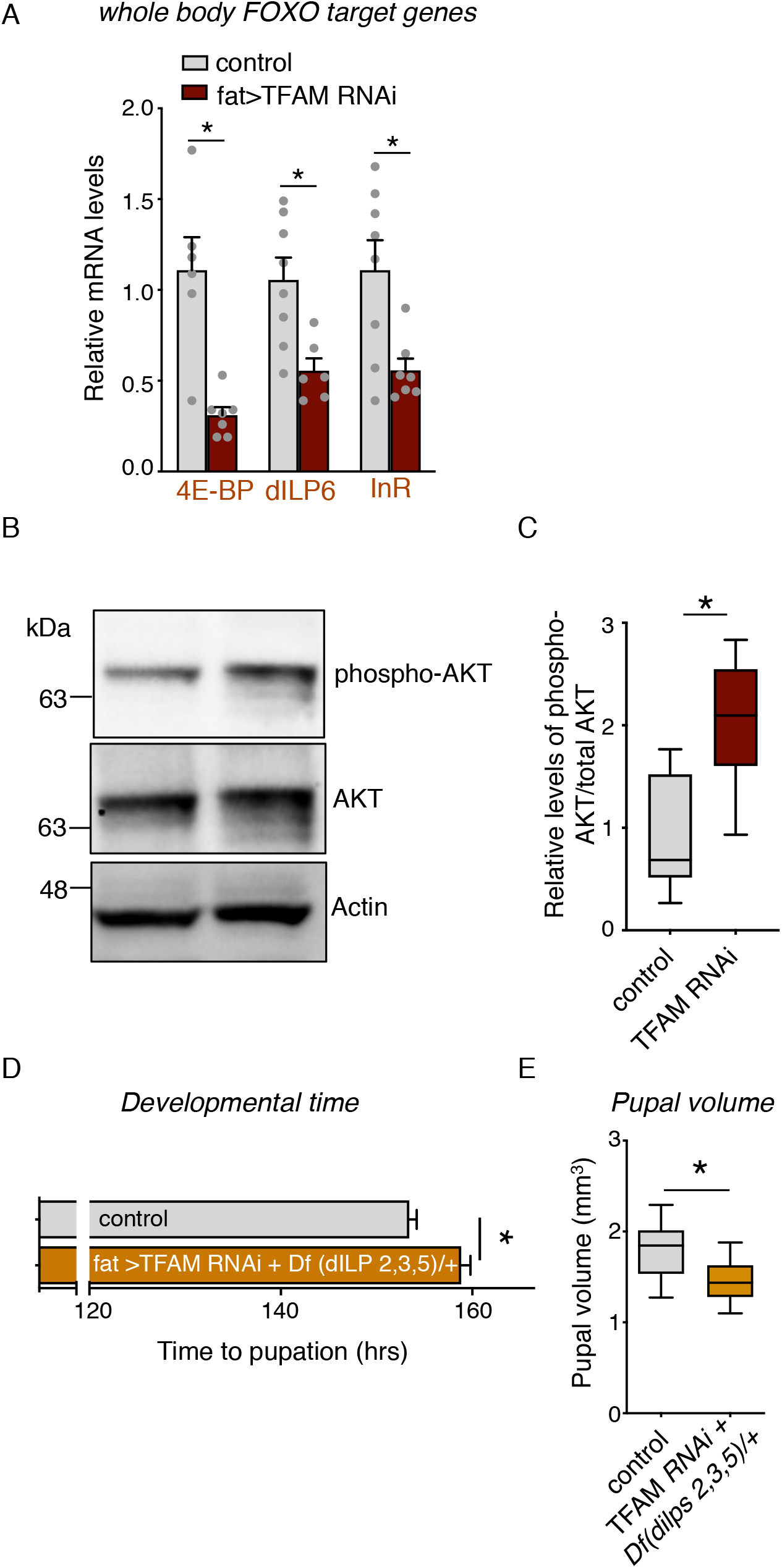
Fat body TFAM knock down increases systemic insulin signaling. (**A**) FOXO transcriptional target genes, 4E-BP, dILP6, and InR were upregulated following TFAM knockdown in the fat body. Total RNA was isolated from control (*r4 > +*) vs TFAM RNAi (*r4 > UAS-TFAM RNAi*) larvae at 96 hrs AEL for qPCR analysis. Data are represented as mean ± SEM with individual data points plotted as symbols (*p < 0.05 unpaired t-test, n = 8 groups per condition). (**B**) Insulin signaling is increased in larvae with fat specific TFAM knock down. Total protein lysates was prepared from control (*r4 > +*) vs TFAM RNAi (*r4 > UAS-TFAM RNAi*) larvae analyzed by western blot analysis. Phospho-Akt, total AKT and actin antibodies were used. (**C**) Quantification of western blots from (B). Data are relative levels of phospho-Akt band intensity corrected for total Akt band intensity. Data are presented as box plots (25%, median and 75% values) with error bars indicating the min and max values (*p < 0.05 unpaired t-test, n > 200 animals). (**D**) Fat body TFAM knockdown does not accelerate growth in animals with reduced insulin signaling. Time to pupation was measured in control (*r4 > +*) larvae vs TFAM RNAi expressed in the fat body (of larvae heterozygous for a deficiency for *dilps 2,3,5* (*r4 > UAS-TFAM RNAi; Df(3L)Ilp2– 3,Ilp53/+*)Data represent mean time to pupation ± SEM (*p < 0.05 Mann-Whitney U test, n > 80 animals per condition) (**E**) Pupal volume was measured in control (*r4 > +*) larvae vs TFAM RNAi expressed in the fat body (of larvae heterozygous for a deficiency for *dilps 2,3,5* (*r4 > UAS-TFAM RNAi; Df(3L)Ilp2–3,Ilp53/+*). Data are presented as box plots (25%, median and 75% values) with error bars indicating the min and max values (*p < 0.05 unpaired t-test, n > 200 animals).

The fat body regulates the IPCs via several secreted adipokines, which can each either activate or inhibit dILP release. We found that the mRNA expression of one inhibitory adipokine, the TNFα homolog, Eiger was significantly reduced in the fat body following TFAM knock down (**Figure 6A**). Suppression of Eiger mRNA levels was also seen following fat body LDH overexpression (**Figure 6B**). We also saw that TFAM knockdown reduced that fat body expression of Impl2, a negative regulator of insulin signaling that functions by binding dILPs and preventing their ability to signal. These results indicate that one way that lowered fat body mitochondrial bioenergetic activity promotes body growth is through reducing the expression of negative regulators of systemic insulin signaling. Indeed, we found that fat specific knockdown of Eiger or its essential processing enzyme, TACE, was sufficient to mimic TFAM knockdown and lead to accelerated growth and development (**Figure 6C and 6D**). Furthermore, we saw that the acceleration in development seen following simultaneous fat body knockdown of TFAM and Eiger, was comparable to the effects of either knockdown alone, suggesting that both function in a similar way (**Figure 6E**).

**Figure 6.**
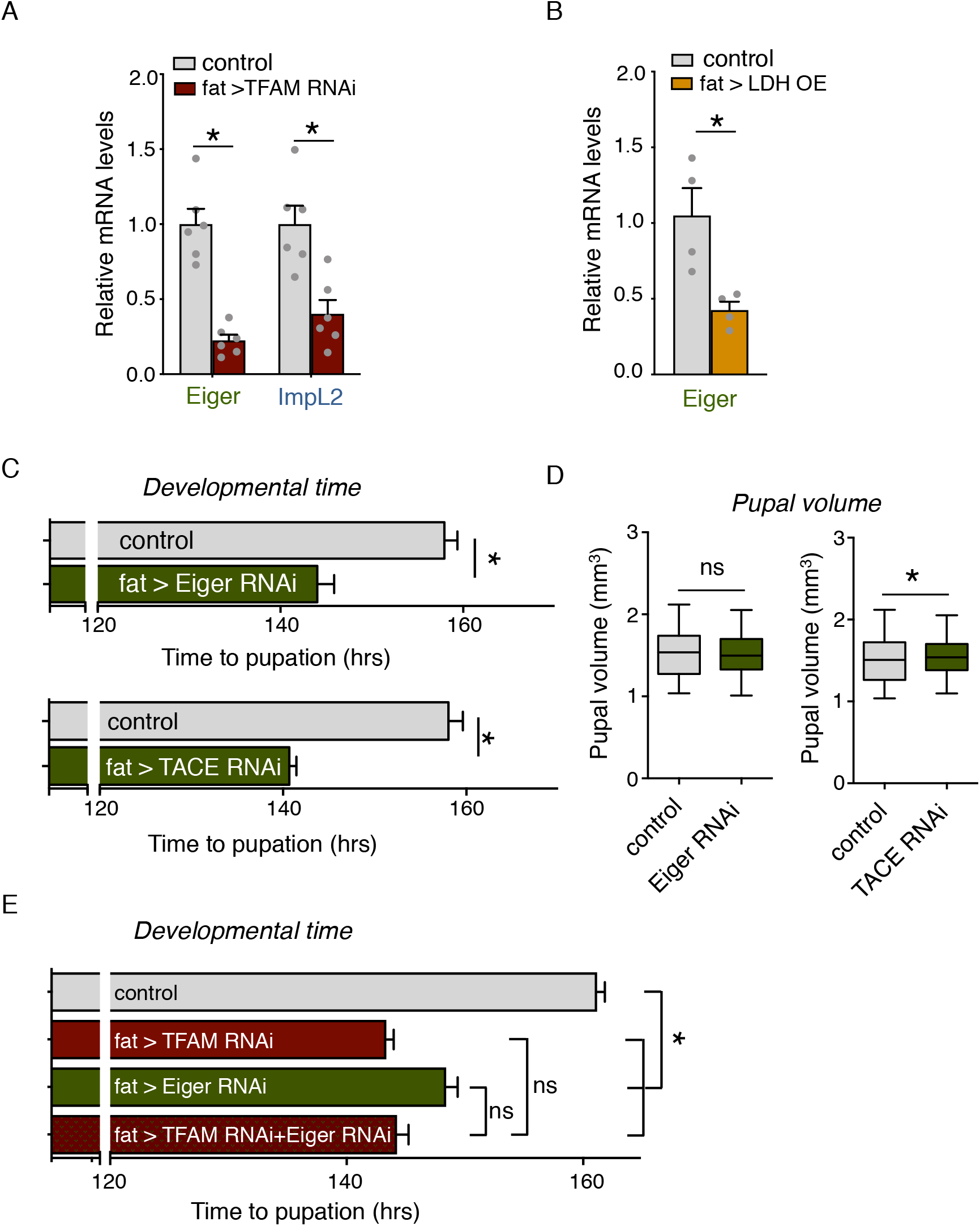
Low bioenergetic fat body suppresses the expression of inflammatory cytokine, TNFα/Eiger to accelerate growth. (**A**) TFAM knock down in fat body reduces the expression of Eiger and Imp-L2, two negative regulators of insulin signaling. Fat bodies were dissected from control (*r4 > +*) vs TFAM RNAi (*r4 > UAS-TFAM RNAi*) larvae at 96 hrs AEL and total RNA was isolated for mRNA-Seq analysis. Data are represented as mean relative levels ± SEM with individual data points plotted as symbols (*p < 0.05 unpaired t-test, n = 6 per condition). (**B**) Fat-specific LDH overexpression reduces Eiger mRNA expression. Total RNA was isolated from control (*r4 > +*) vs LDH overexpressing (OE) (*r4 > UAS-LDH*) larvae at 96 hrs AEL and analyzed by qRT-PCR. Data are represented as mean relative levels ± SEM with individual data points plotted as symbols (*p < 0.05 unpaired t-test, n = 4 per condition). (**C**) Fat body specific knockdown of Eiger or its processing enzyme, TACE accelerates larval development. Time to pupation was measured in control (*r4 > +*) vs Eiger RNAi (*r4 > UAS-Eiger RNAi*) or TACE RNAi (*r4 > UAS-TACE RNAi*) larvae. Data represent mean time to pupation ± SEM (*p < 0.05 Mann-Whitney U test, n > 80 animals per condition) (**D**) Fat knock down of Eiger has no effect on pupal size while TACE knock down lead to a slight increase in pupal size compared to controls. Data are presented as box plots (25%, median and 75% values) with error bars indicating the min and max values (*p < 0.05, ns = not significant, unpaired t-test, n > 200 animals) (**E**) Fat body knockdown of TFAM and Eiger RNAi, both alone and together leads to accelerated larval development. Time to pupation was measured in control (*r4 > +*) larvae vs larvae expressing Eiger RNAi (*r4 > UAS-Eiger RNAi*), TACE RNAi (*r4 > UAS-TACE RNAi*), or both Eiger and TACE RNAi. Data represent mean time to pupation ± SEM (*p < 0.05, ns = not significant, Mann-Whitney U test, n > 100 animals per condition).

## Discussion

### Mitochondrial metabolism as a driver of animal growth and development

Our central finding is that the regulation of fat body mitochondrial metabolism can drive whole body growth and development **(Figure 7)**. In recent years, there has been a resurgence of interest in the links between metabolism and animal development. This interest stems not just from the fact that metabolism provides the molecular building blocks needed for tissue and body growth, but because it is also becoming clear that specific metabolic processes can actively drive changes in cell function and behavior (Drummond-Barbosa and Tennessen, 2020; Gandara and Wappner, 2018; Miyazawa and Aulehla, 2018; Sieber and Spradling, 2017). Mitochondria, in particular, play a central role in this metabolic control of development. For example, in *Drosophila* reprogramming of mitochondrial function has been shown to direct changes in oogenesis (Sieber et al., 2016), stem cell behavior (Homem et al., 2015; Schell et al., 2017; Senos Demarco et al., 2019), cellular differentiation (Mitra et al., 2012), morphogenesis (Chowdhary et al., 2020), and growth (Noguchi et al., 2011). In each of these cases mitochondria function cell-autonomously to mediate their effects. Our results add to this work by showing how alterations in mitochondria in one tissue can induce whole-body effects on animal development. We found that not only did fat body mitochondrial morphology and bioenergetic activity change during larval development and in response to nutrient availability, but that lowering mitochondrial bioenergetic activity alone in the fat body was enough to accelerate larval growth. These findings place adipose mitochondrial metabolism as a central regulator of nutrient-dependent larval development. Moreover, given the functional similarity of the fat body to vertebrate adipose tissue and liver, we propose that mitochondrial reprogramming in these nutrient-sensing tissues may also control whole-body physiology and growth.

**Figure 7.**
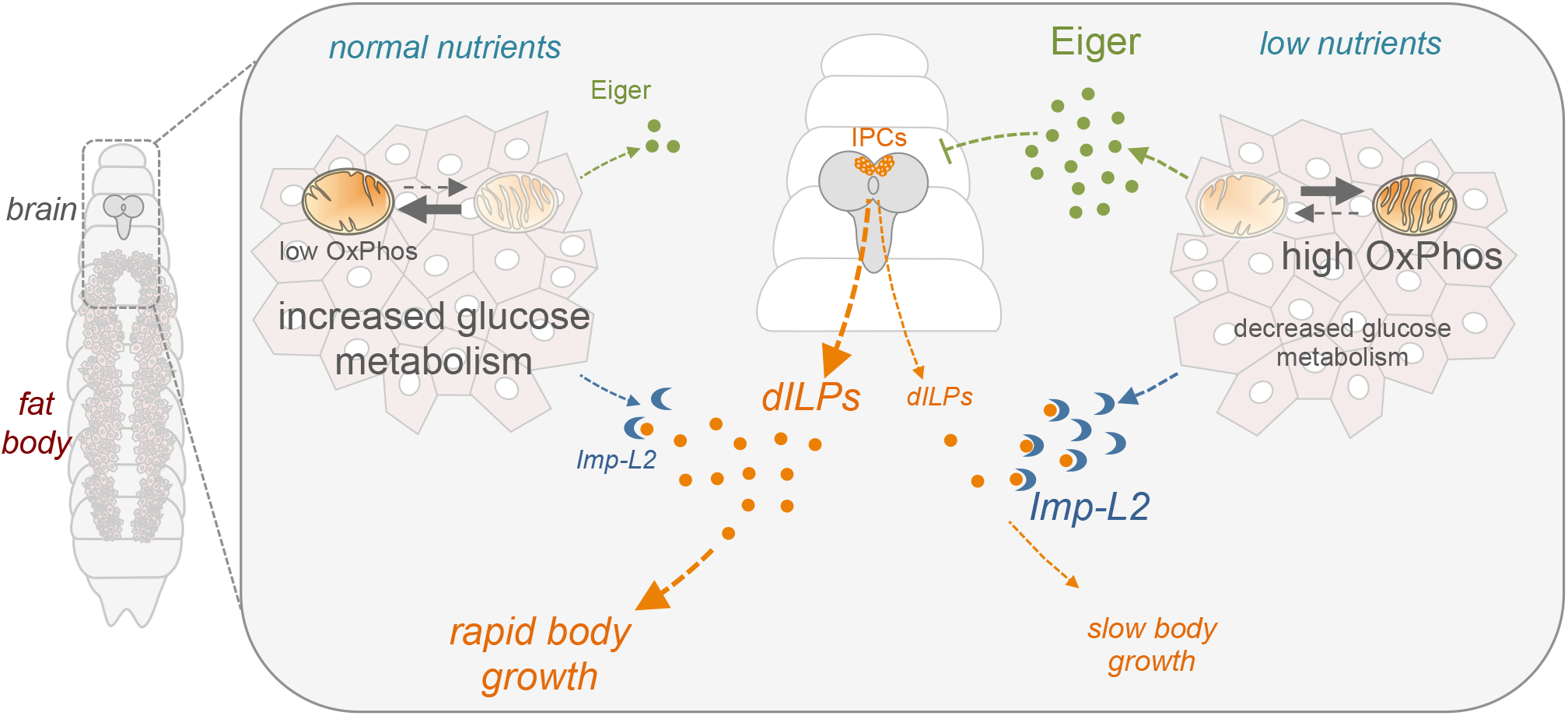
Adipose mitochondrial metabolism controls body growth by modulating cytokine and insulin signaling. A model for how fat body mitochondrial metabolism can control whole-body growth. When nutrients are abundant fat body mitochondria engage in low OxPhos activity, and as a result, fat body glucose metabolism is increased, and the expression of Eiger and Imp-L2, two negative regulators of insulin signaling, is reduced. Under these conditions, dILPs secreted from the brain IPC cells can then promote systemic insulin signaling to drive rapid body growth and development. In contrast, when nutrients are limiting fat body mitochondria increase their OxPhos activity, glucose metabolism is reduced, and the expression of both Eiger and Imp-L2 is elevated. This then leads to suppressed systemic insulin signaling and slower growth and development. TFAM knockdown reduces mitochondrial OxPhos activity and thus can enhance the rapid growth in rich nutrients and reverse the slow growth in low nutrients.

### Fat body mitochondria modulate systemic insulin signaling

We discovered that fat body mitochondrial metabolism exerts its organismal growth effects through non-autonomous control of whole-body insulin signaling. Here there are parallels with mouse studies linking adipose mitochondria with insulin sensitivity in the context of diabetes and obesity. These links appear bidirectional, and they vary depending on the tissues studied, the experimental models used, and the features of mitochondrial function tested (Koliaki and Roden, 2014, 2016). For example, adipose-specific deletion of TFAM increases mitochondrial oxidation and can protect mice against obesity and insulin resistance (Vernochet et al., 2012). However, another study showed that inactivation of TFAM in adipose tissue results in decreased complex I and IV activity, which is similar to changes observed in skeletal muscle following tissue-specific TFAM knock out (Wredenberg et al., 2002). Moreover, inherited or acquired mitochondrial abnormalities can lead to lower mitochondrial function resulting in insulin resistance and type 2 diabetes mellitus (Kelley et al., 2002; Koliaki and Roden, 2016; Shulman, 2004; Szendroedi et al., 2011). Deregulation of growth signaling pathways, such as insulin signaling, can also impair mitochondrial function (Hu et al., 2021). Our results suggest that, in addition to these pathological contexts, adipose mitochondria can function to control normal systemic physiology and growth.

### Mitochondrial metabolism as a regulator of adipose cytokine production

The fat body plays a central role in coupling changes in dietary nutrients to the regulation of systemic insulin and body growth. This role is mediated in large part through adipokines that signal from the fat body to the brain to control the release of dILPs (Texada et al., 2020). We found that changes in expression of one adipokine, Eiger, were important for the effects of fat body mitochondria on systemic insulin signaling and growth. Eiger is the *Drosophila* homolog of the inflammatory cytokine TNFα, and can control diverse functions including cell proliferation, differentiation, immunity, and death (Igaki and Miura, 2014). In *Drosophila* larvae growing in low nutrient conditions, Eiger is released by the fat body into the hemolymph where it signals to the brain IPCs to suppress the expression of dILPs and, as a result, reduce body growth (Agrawal et al., 2016). These effects rely on suppression of TOR signaling in the fat body. Our findings raise the possibility that changes in mitochondrial metabolism couple TOR signaling to the regulation of Eiger. Indeed, suppression of Eiger expression occurred in response to lowering bioenergetic activity, a metabolic state that we see in fat bodies from larvae grown in nutrient rich conditions when TOR activity is high. Furthermore, regulation of mitochondrial activity is a conserved downstream function of TOR signaling (Morita et al., 2013), particularly in response to changes in mRNA translation, which has been shown to regulate fat body control of systemic insulin and growth (Delanoue et al., 2010; Marshall et al., 2012; Rideout et al., 2012). Interestingly, in mammals, TNFα is also produced from adipose tissue in response to different nutrient signals (Hotamisligil et al., 1995; Hotamisligil et al., 1993; Hotamisligil and Spiegelman, 1994). Thus, it is interesting to speculate whether these effects may also be triggered by alterations in fat mitochondrial metabolism.

In *Drosophila*, TNFα exerts many of its different effects through paracrine and endocrine signaling, often in response to extracellular stress cues such as tissue damage or infection (Igaki and Miura, 2014; Mabery and Schneider, 2010; Parisi et al., 2014; Sanchez et al., 2019). Our findings raise the possibility that in these situations, the induction of Eiger may also be triggered by alterations in mitochondrial function. Interestingly, some of the downstream effects of Eiger signaling have been shown to require alterations in mitochondrial metabolism, suggesting that links between mitochondria and Eiger may be bi-directional (Kanda et al., 2011; Keller et al., 2011).

We also found that fat body TFAM knockdown reduced expression of Imp-L2, the fly homolog of the mammalian insulin growth factor binding proteins (Honegger et al., 2008). Secreted Imp-L2 can bind to circulating dILPs and prevent their ability to signal. Thus, lowered fat body mitochondrial bioenergetics appears to enhance growth by suppressing two secreted factors, Eiger and Imp-L2, which are negative regulators of insulin signaling. Indeed, Imp-L2 mutants show accelerated development, particularly in low nutrient conditions (Sarraf-Zadeh et al., 2013). Interestingly, in the context of lifespan regulation, production of Imp-L2 from adult muscle is increased by deregulation of mitochondrial function (Owusu-Ansah et al., 2013), suggesting that a mitochondria-Imp-L2 link may operate in different tissues and in different physiological contexts.

### Fat body mitochondria and metabolic reprogramming

We found that lowered fat body mitochondrial bioenergetics led to the reprogramming of glucose metabolism through upregulation of glucose metabolism gene expression. Moreover, these mitochondrial and glucose metabolic changes altered eiger gene expression to control systemic insulin and body growth. How might these changes in fat cell metabolism occur and how might they lead to altered gene expression? One intriguing possibility is that they rely on changes in TCA cycle activity and TCA metabolites. For example, recent studies in mammalian cells have shown how TCA cycle intermediates such as alpha-ketoglutarate, succinate and fumarate can regulate histone acetylation and other epigenetic modifications to control gene expression (Baksh and Finley, 2021; Martinez-Reyes and Chandel, 2020). Hence, when we lower fat body OxPhos by TFAM knockdown, these intermediates may trigger increased glucose metabolism gene expression. Similar mechanisms may also mediate the effects on Eiger gene expression. Here the situation may be analogous to activated macrophages, which switch from OxPhos to glycolysis to generate ATP, and then use their TCA cycle to produce succinate, which controls cytokine gene expression (Mills et al., 2016; Ryan et al., 2019; Ryan and O’Neill, 2020; Tannahill et al., 2013). Lowering mitochondrial bioenergetics (by TFAM knockdown) and the subsequent alterations in glucose metabolism may also lead to rewiring and altered fueling of the TCA cycle. For example, human cells with mitochondrial DNA mutations and OxPhos defects drive their TCA cycle using glutamine instead of glucose (Chen et al., 2018). In addition, increased glucose and glutamine availability in mouse pancreatic islet cells has been shown to rewire TCA cycle metabolism in order to promote insulin secretion (Zhang et al., 2021). Interestingly, gene ontology analyses of our fat body RNA seq data showed enrichment in glutamate and glutamine metabolism among the genes upregulated following TFAM knockdown.

Our results underscore the importance of mitochondrial and glucose metabolic plasticity in the control of animal development. This developmental metabolic plasticity is emphasized by other studies in *Drosophila*. For example, reorganization of mitochondrial and glucose metabolism occurs across embryo to larval development to support organismal growth (Matoo et al., 2019; Tennessen et al., 2011; Tennessen et al., 2014). In addition, genetic loss of LDH leads to altered flux through other glucose metabolic pathways to maintain growth (Li et al., 2019). Finally, mitochondrial pyruvate import has been shown to be dispensable for homeostasis in normal food, but absolutely required for survival on a carbohydrate-only diet (Bricker et al., 2012).

In summary, our findings show how altering mitochondrial activity in a key nutrient-sensing tissue - the fat body – can drive overall body growth. Given that the fat body plays a central role as both a sensor of different environmental cues (such as hypoxia, infection and oxidative stress), and as a regulator of other organismal responses such as immunity, fecundity and lifespan, it will be interesting to explore whether the mitochondrial mechanisms we identified may also operate in these contexts.

## Materials and Methods

### Drosophila Strains

The following strains were used: *w*^*1118*^, *r4-GAL4, da-GAL4, (*Bloomington Drosophila Stock Center (BDSC), *UAS-dLDH (Li et al., 2017)* (Gift from Jason Tennessen) *UAS-TFAM RNAi #1 (VDRC 37819), UAS-MPC1 RNAi (VDRC 15858), UAS-Eiger RNAi (VDRC 108814), UAS-TACE RNAi (VDRC 2733)* (Vienna Drosophila RNAi Center), *UAS-TFAM RNAi #2 (4217R-2), UAS-TFAM RNAi #3 (4217R-3)* (Fly Stocks of National Institute of Genetics - NIG-FLY).

### *Drosophila* food and genetics

Flies were raised on standard medium containing 150 g agar, 1600 g cornmeal, 770 g Torula yeast, 675 g sucrose, 2340 g D-glucose, 240 ml acid mixture (propionic acid/phosphoric acid) per 34 L water and maintained at 25 °C. For all GAL4/UAS experiments, homozygous GAL4 lines were crossed to the relevant UAS line(s) and the larval progeny were analyzed. Control animals were obtained by crossing the relevant homozygous GAL4 line to flies of the same genetic background as the particular experimental UAS transgene line.

### MitoTracker Red staining

Fat bodies from 96hrs AEL larvae were inverted in cold PBS and stained with MitoTracker Deep Red FM (1: 1000 dilution of 1 mM, Molecular probes M22426) for 40 mins and fixed at room temperature using 8% PFA for 30 mins. After washing three times, mounted using Vecta Shield mounting medium. The mitochondrial images were acquired through Zeiss confocal microscope LSM 880.

### LysoTracker staining

Fat bodies from 96hrs AEL larvae were dissected and incubated in LysoTracker (1:1000, Thermofisher Scientific, L7528) for 10 mins on glass slides at room temperature. Fat bodies were then immediately imaged using a Zeiss Observer Z1 microscope using Axiovision software.

### Lipid droplet staining with BODIPY

Fat bodies from 96hrs AEL larvae were inverted in cold PBS and fixed at room temperature using 8% PFA for 30 mins. Tissues were then incubated in BODIPY for 30 minutes. After washing three times, fat bodies were dissected and mounted using Vecta Shield mounting medium. The fat body images were acquired through Zeiss confocal microscope LSM 880.

### Glucose uptake measurement

The fluorescently labeled glucose analogue, 2-deoxy-2-[(7-nitro-2,1,3-benzoxadiazol-4-yl)amino]- D-glucose (2-NBDG) (Invitrogen N13195) was used as a probe for monitoring glucose uptake. Dissected larval fat tissues were incubated with 2-NBDG at 300 uM concentration in PBS for 30 mins at room temperature. Then, the samples were washed with PBS and fixed in 4% PFA-PBS for 30 mins at room temperature. After washing there times with PBS, fat tissue samples were mounted using Vecta Shield mounting medium. The fat tissue images were acquired through Zeiss confocal microscope LSM 880.

### Glycogen staining with periodic acid solution (PAS)

For staining of glycogen, 96 hrs. AEL larval fat bodies were dissected in PBS, fixed using 4% paraformaldehyde in PBS for 20 min, and washed twice with 1% BSA in PBS (Yamada et al., 2018). Samples were incubated with periodic acid solution (Merck) for 5 min and washed twice with 1% BSA in PBS. Fat bodies were then stained with Schiff’s reagent (Merck) for 15 min, washed twice, and mounted in 50% glycerol in PBS. Images were acquired with a Zeiss Observer. Z1. Images were analyzed using NIH Image J software by measuring the average intensity of staining in 10 different random regions per tissue sample, a total of 15 to 20 fat tissues were analyzed in each condition.

### Transmission Electron Microscopy (TEM)

Fat bodies were dissected from different stages of larvae and fixed with 2% paraformaldehyde and 2.5% glutaraldehyde in 0.1 M cacodylate buffer, pH 7.4 for at least 2 hours. After washing three times with the same buffer, supernatant was discarded and warm 1% agarose was added over the pellet. After the agarose has cooled, the embedded pellet was cut into pieces. The pieces were post-fixed in 1% osmium tetroxide in cacodylate buffer for 1 hour, dehydrated in a water/ acetone series and embedded in Epon resin. Ultrathin sections (~60 nm thickness) were cut in a Leica EM UC7 ultra microtome using a diamond knife and stained with 2% aqueous uranyl acetate and Reynolds’s lead citrate. The sections were examined in a Hitachi H7650 transmission electron microscope at 80 kV. The images were acquired through an AMT 16000 CCD mounted on the microscope. TEM images were analyzed using NIH Image J software by measuring the mitochondrial cross-sectional area and quantifying the inner membrane junctions.

### Mitochondrial isolation from *Drosophila* larvae

Larvae from 96 hrs. AEL (50 larvae per group) were frozen on dry ice or stored −80°C freezer. Larvae were thawed on ice and lysed in 500 ul of STE+ BSA buffer by homogenization. Then the lysate was spun at 600 g at 4°C for 10 minutes. Supernatant was collected and centrifuged at 7 000 g for 15 min at 4°C. Next, supernatant was discarded and the pellet was washed in ice-cold STE buffer (250 mM sucrose, 10 mM Tris-HCl pH 7.0 and 0.2 mM EDTA pH 8.0) and resuspended in 50 ul STE buffer. The pellet containing mitochondria were aliquoted and frozen in dry ice first and transferred to – 80°C freezer for future complex activity assays. Mitochondrial protein concentration was determined using the Dc-protein determination kit (BioRad).

### Measurement of mitochondrial citrate synthase and complex IV activity

For all the enzymatic assays, frozen isolated mitochondria were thawed on ice, resuspended in STE buffer at 1 mg/ml and permeabilized with n-Dodecyl **β**-D-maltoside at 0.015% final concentration (Sriskanthadevan et al., 2015). All the enzymatic assays consist of 5 – 10 μg of mitochondrial protein. Citrate synthase activity was measured based on the chemical coupling of CoASH, released from acetyl-CoA during the enzymatic synthesis of citrate to DTNB (Ellman’s reagent, 5,5’-dithiobis(2-nitrobenzoic acid), and the release of the absorbing mercaptide ion was monitored at 412 nm (Kaplan and Colowick, 1955). The reaction mixture contained 5 μg of mitochondrial protein, 0.4 mg/ml DTNB, 0.3 mM acetyl Co-A in 0.1 M Tris-HCl pH 8.0 buffer. The reaction was started by the addition of 5 mM oxaloacetic acid. An extinction coefficient of 13.6 mM-1 cm-1 was used to calculate activities. Complex IV activity was measured by oxidation of ferrocytochrome c (Trounce et al., 1996). Ferrocytochrome c was prepared by reducing cytochrome c with sodium ascorbate followed by dialysis for 24 hours (Zheng et al., 1989).

### Measurement of *Drosophila* developmental time and pupal volume

For measuring development timing to pupal stage, newly hatched larvae were collected at 24 hrs. AEL and placed in food vials (50 larvae per vial). The number of pupae was counted twice each day. For each experimental condition, a minimum of five replicates was used to calculate the mean percentage of pupae per time point. For pupal volume analysis, pupae were imaged using a Zeiss Discovery V8 Stereomicroscope with Axiovision imaging software. Pupal length and width were measured and pupal volume was calculated using the formula, volume=4/3π(L/2)(l/2)2.

### Pupal imaging

Pupal images were obtained using a Zeiss Stereo Discovery V8 microscope using Axiovision software. Microscopy and image capture was performed at room temperature and captured images were processed using Photoshop CS5 (Adobe).

### Preparation of larval protein extracts

*Drosophila* larvae (96 hrs. AEL) were lysed with homogenization and sonication in a buffer containing 20 mM Tris-HCl (pH 8.0), 137 mM NaCl, 1 mM EDTA, 25% glycerol, 1% NP-40 and with following inhibitors 50 mM NaF, 1 mM PMSF, 1 mM DTT, 5 mM sodium ortho vanadate (Na_3_VO_4_) and Protease Inhibitor cocktail (Roche Cat. No. 04693124001) and Phosphatase inhibitor (Roche Cat. No. 04906845001), according to the manufacturer instructions.

### Western blots, immunostaining and antibodies

Protein concentrations were measured using the Bio-Rad Dc Protein Assay kit II (5000112). Protein lysates (60 μg) were resolved by SDS-PAGE and electro transferred to a nitrocellulose membrane, subjected to western blot analysis with specific antibodies, and visualized by chemiluminescence (enhanced ECL solution (Perkin Elmer)). Primary antibodies used in this study were: anti-phospho-AKT-Ser505 (1:1000, Cell Signalling Technology #4054), anti-AKT (1:1000, Cell Signaling Technology #9272), anti-actin (1:1000, Santa Cruz Biotechnology, # sc-8432). Goat secondary antibodies were purchased from Santa Cruz Biotechnology (sc-2030, 2005, 2020)

### mRNA Seq analysis

RNA-sequencing was conducted by the University of Calgary Centre for Health Genomics and Informatics. The RNA Integrity Number (RIN) was determined for each RNA sample (6 replicates per each condition were used). Samples with a RIN score higher than 8 were considered good quality, and Poly-A mRNA-seq libraries from such samples were prepared using the Ultra II Directional RNA Library kit (New England BioLabs) according the manufacturer’s instructions. Libraries were then quantified using the Kapa qPCR Library Quantitation kit (Roche) according to the manufacturer’s directions. Finally, RNA libraries were sequenced for 100 cycles using the NovaSeq 6000 Sequencing System (Illumina).

### Quantitative RT-PCR measurements

Total RNA was extracted from larvae using TRIzol according to manufacturer’s instructions (Invitrogen; 15596-018). RNA samples isolated from same number of larvae (control vs experimental) were DNase treated (Ambion; 2238G) and reverse transcribed using Superscript II (Invitrogen; 100004925). The generated cDNA was used as a template to perform qRT–PCRs (ABI 7500 real time PCR system using SyBr Green PCR mix) using gene-specific primers. PCR data were normalized to RpL32. All primer sequences are listed in supplemental table 1.

### Statistical analysis

All qRT-PCR data and quantification of immunostaining data were analyzed by Students t-test, two-way ANOVA followed by post-hoc students t-test, or Mann-Whitney U test where appropriate. All statistical analysis and data plots were performed using Prism statistical software. Differences were considered significant when p values were less than 0.05.

## Acknowledgements

We thank Jason Tennessen for the gift of fly stocks. Stocks obtained from the VDRC, the NIG-Fly Stock Centre, Kyoto, Japan and the Bloomington Drosophila Stock Center (NIH P40OD018537) was used in this study. We thank Wei-Xiang Dong for technical support with electron microscopy and Qingrun Zhange for bioinformatic support. This work was supported by a CIHR project grant and a Cancer Research Society grant to S.S.G. M.J.T. was supported by an NSERC summer studentship.

---

**Figure S1 (related to Figure 2).**
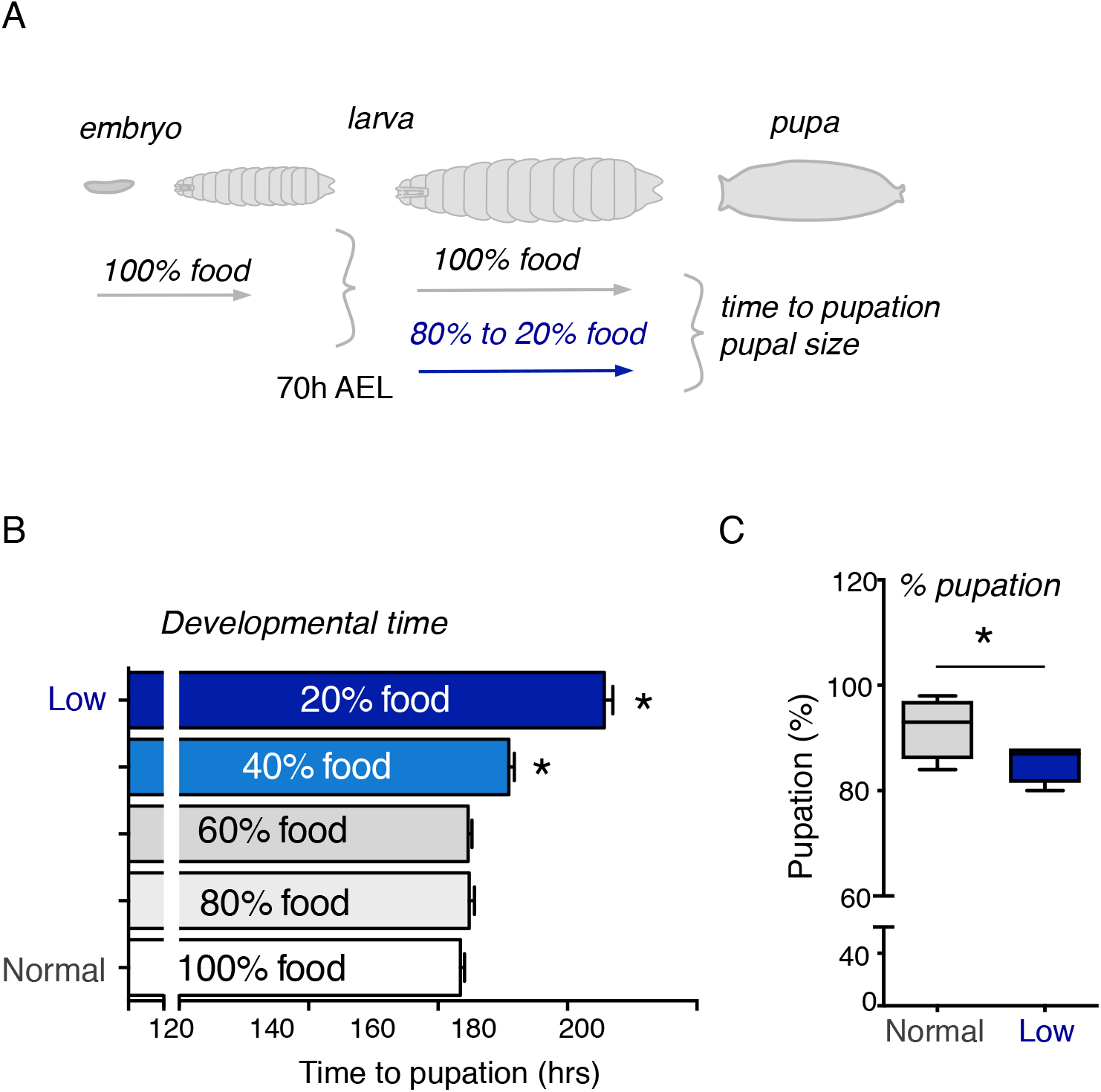
Fat body mitochondrial architecture altered in larvae grown in low nutrient condition. **(A)** A schematic is shown depicting the larval growth phase of development and the experimental setup. (**B**) Larvae grown on normal food were transferred to diluted food conditions (80% to 20% of the normal food) at 70 hrs AEL and the time to pupation was measured. (**C**) Larvae grown on normal food were transferred to diluted food conditions (20% food) at 70 hrs AEL and the % of pupation was measured.

**Figure S2 (related to Figure 2).**
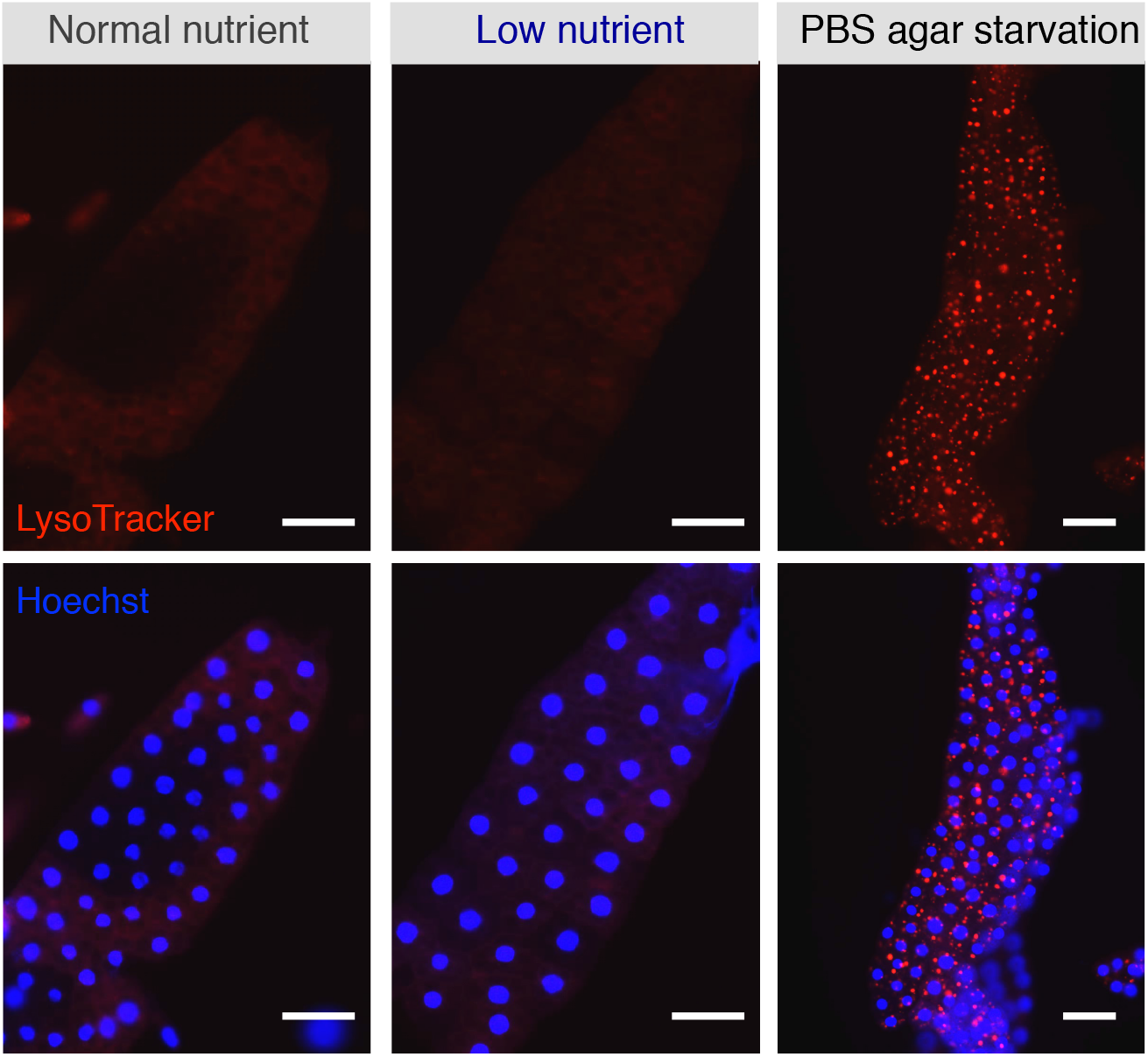
*Drosophila* larvae grown on low nutrient condition does not induce autophagy in fat body. Animals from embryo to 70 hrs (AEL) maintained in normal food and then transferred to normal food, low nutrient food (diluted 20% of normal food) or starved on PBS for 24 hrs. Fat bodies were then dissected, stained with LysoTracker and then imaged. Red indicates LysoTracker stain and blue indicates Hoechst DNA stain. Scale bar = 50 μm.

**Figure S3 (related to Figure 2).**
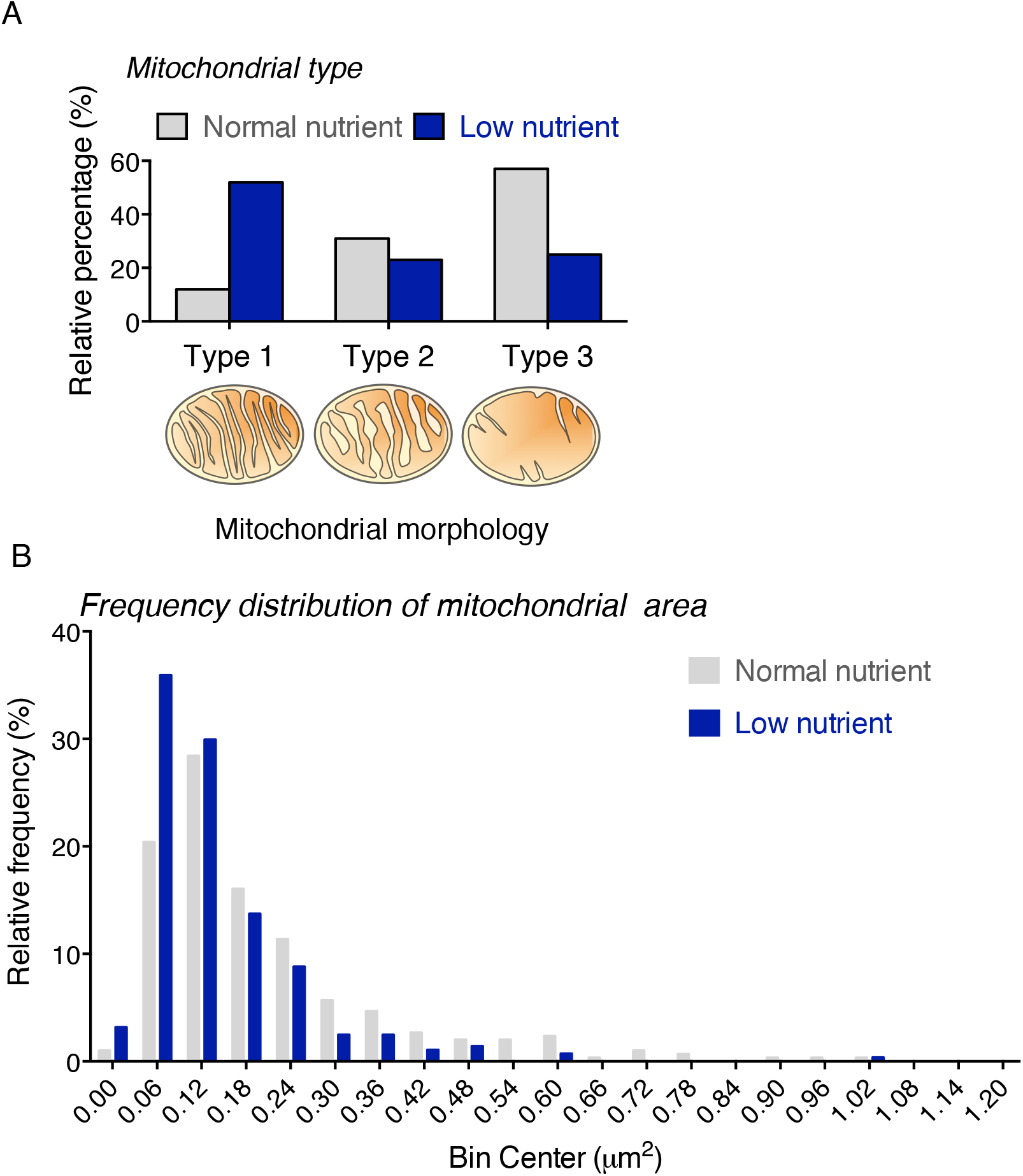
Fat body mitochondrial architecture altered in larvae grown in low nutrient condition. **(A**) Analysis of TEM images of larvae grown in normal and low nutrient food. Percentage of mitochondrial type were quantified by counting the number of mitochondria from 20 to 25 different TEM images taken in normal and low nutrient conditions. Low nutrient fat body consists of highest percentage of type 1 mitochondria (similar to highly respiring muscle mitochondria) compare to normal food which consists of type III mitochondria. (**B**) Frequency distribution histogram of mitochondrial cross- sectional area from fat body TEM images. Larvae treated with low nutrient for 24 hrs has lower mitochondrial cross-sectional area compared to the control larvae.

**Figure S4 (related to Figure 3).**
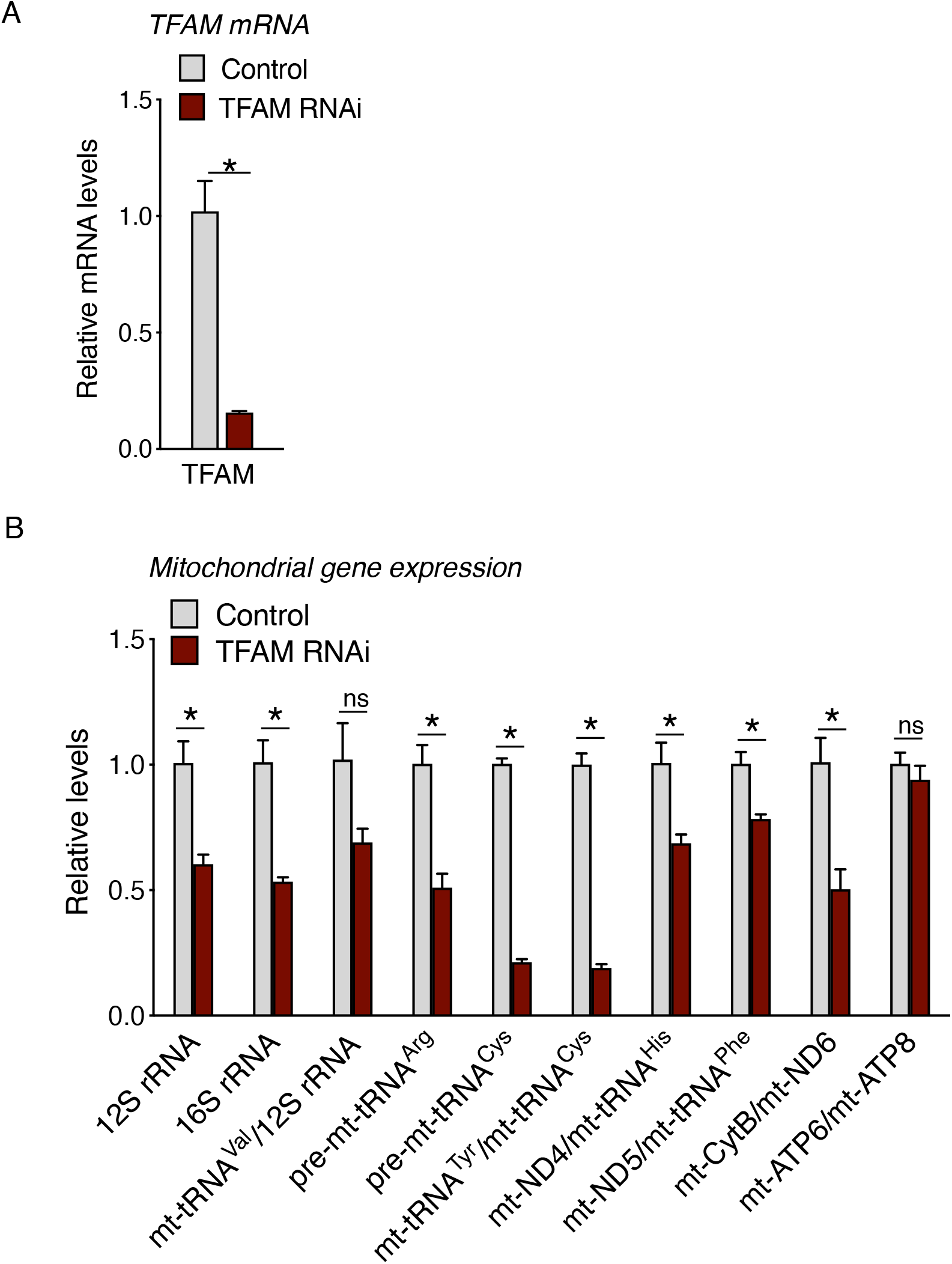
Whole body TFAM knockdown (RNAi # 1) leads to reduced mitochondrial gene expression. (**A**) TFAM expression reduced significantly in TFAM knock down animals. (**B**) Expression of mitochondrial DNA encoded genes in TFAM knock down animals. Data represented as mean ± SEM (* p < 0.05 unpaired t-test, ns = not significant.)

**Figure S5 (related to Figure 3):**
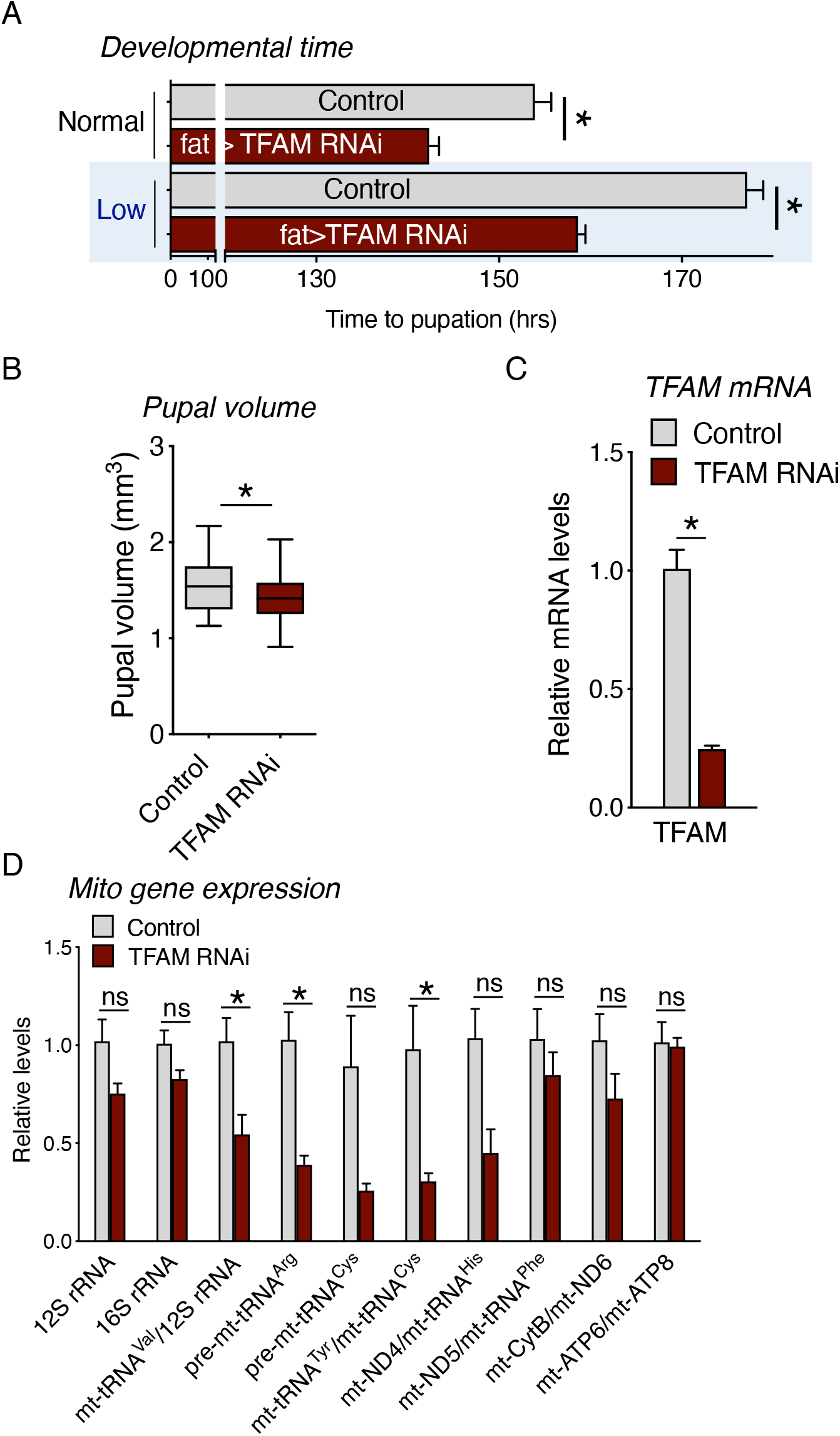
(**A-B**) TFAM knockdown (RNAi # 2) in fat body speeds up animal growth and development. (**C**) TFAM expression reduced significantly in TFAM knock down animals. (**D**) Expression of mitochondrial DNA encoded genes in TFAM knock down animals. Data represented as mean ± SEM (* p < 0.05 unpaired t-test, ns =

**Figure S6 (related to Figure 3):**
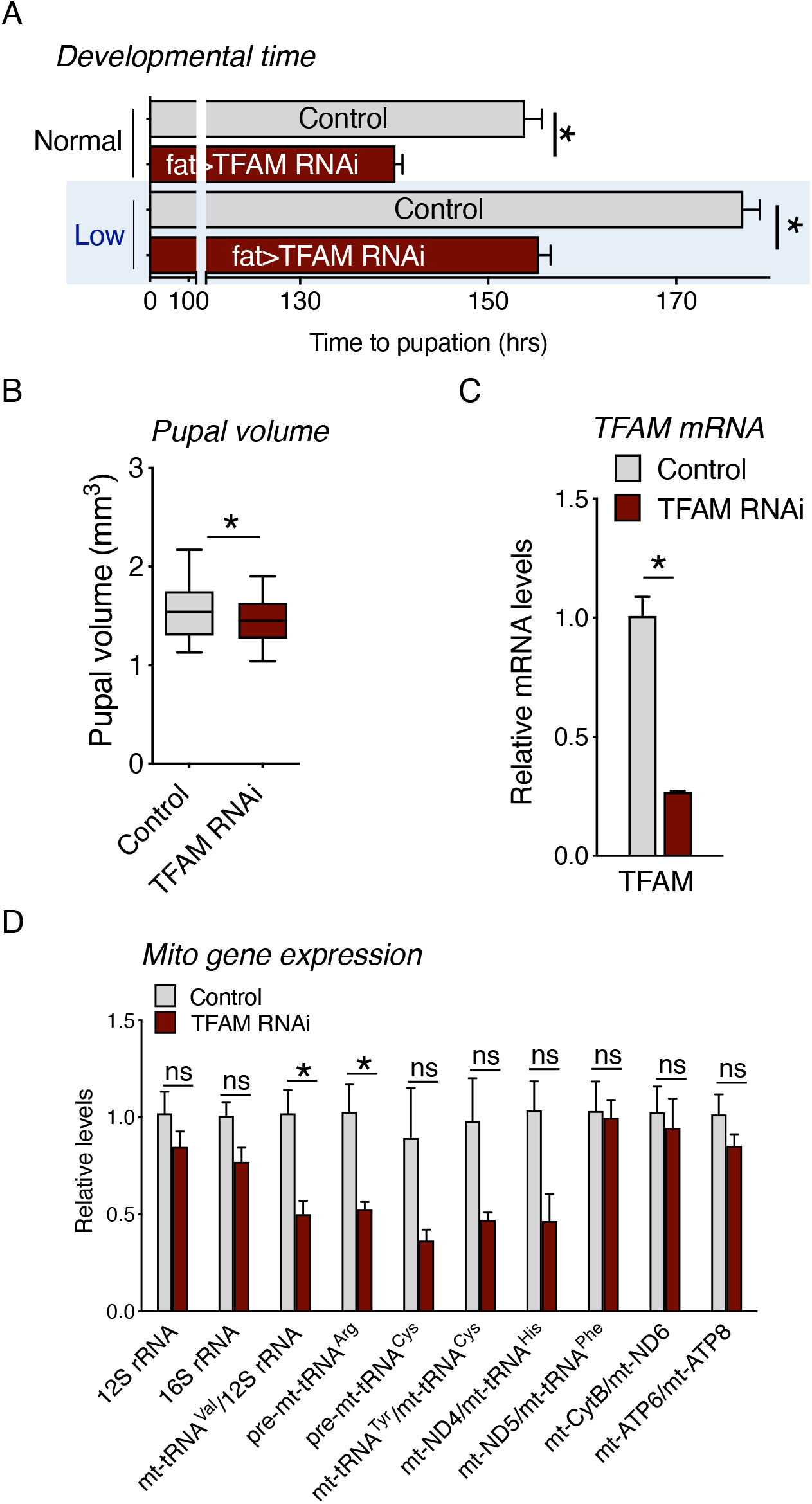
(**A-B**) TFAM knockdown (RNAi # 3) in fat body speeds up animal growth and development. (**C**) TFAM expression reduced significantly in TFAM knock down animals. (**D**) Expression of mitochondrial DNA encoded genes in TFAM

**Figure S7 (related to Figure 3).**
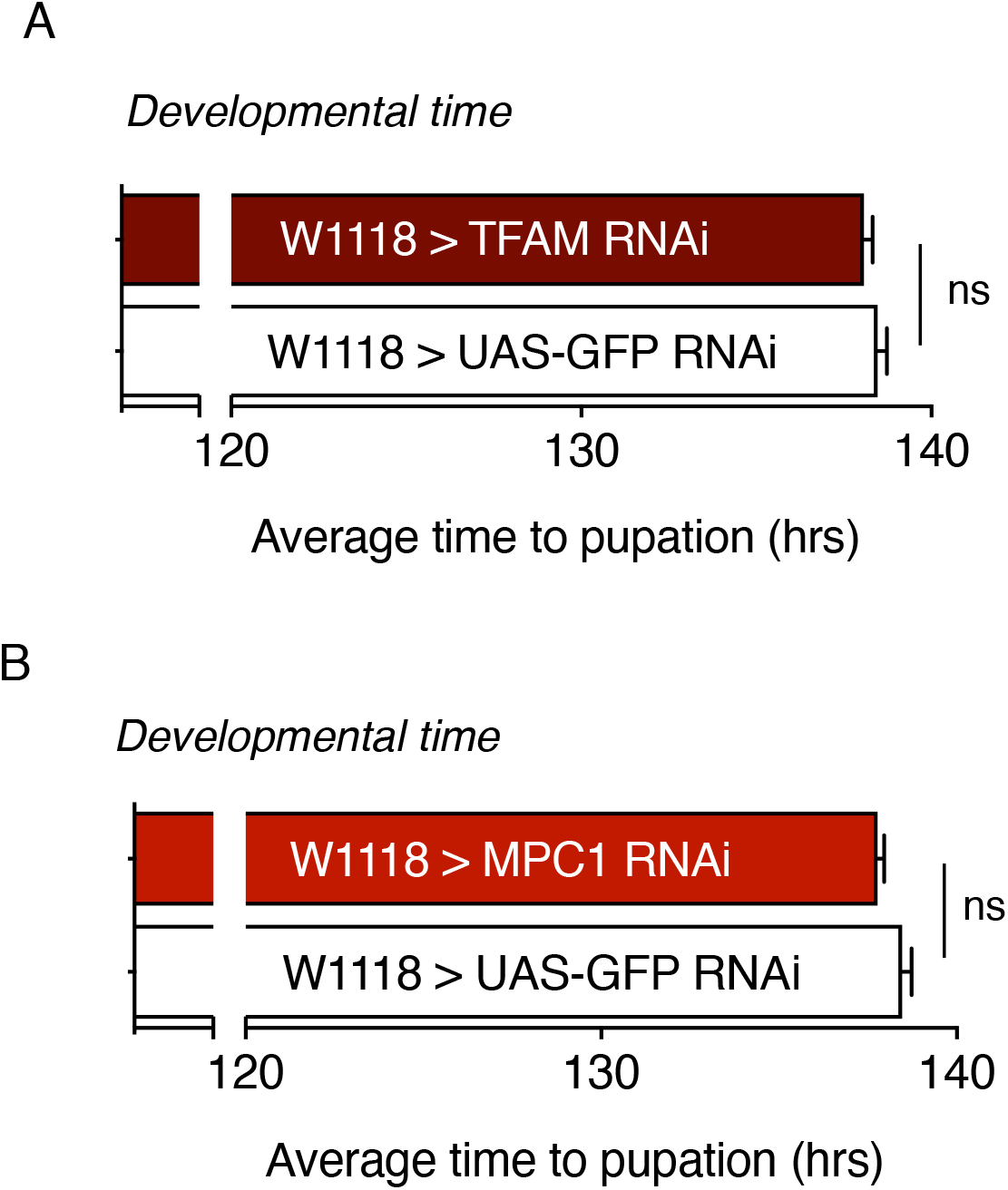
No driver controls for TFAM RNAi (RNAi #1) and MPC1 RNAi show no change in the average time to pupation. n > 400 animals. Data represented as mean ± SEM (* p < 0.05 unpaired t-test, ns = not significant.)

**Figure S8 (related to Figure 4).**
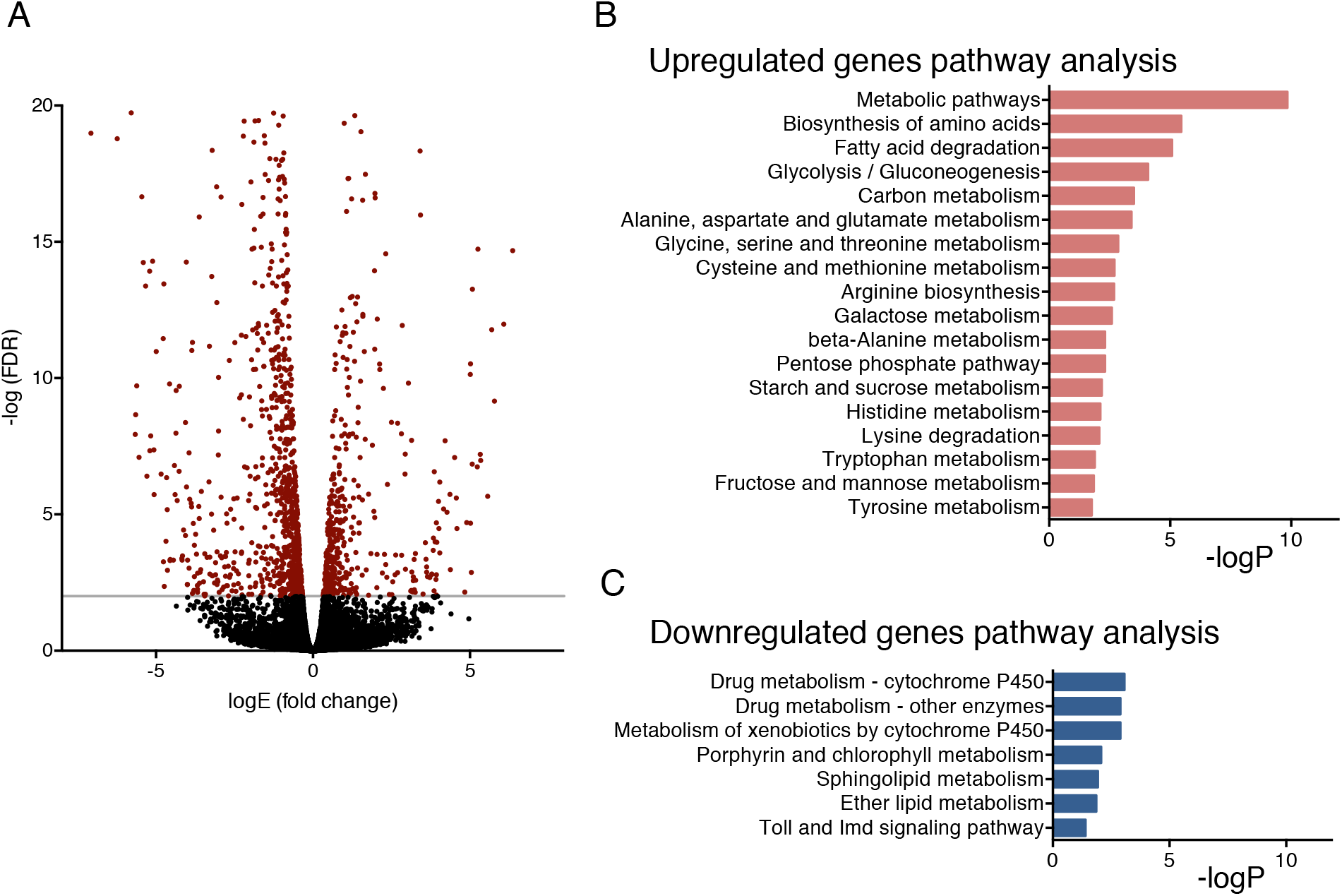
TFAM knock down fat body mRNA Seq Analysis. (**A**) Volcano plot of fat body TFAM knock down mRNA Seq differentially expressed genes (DEGs). (**B-C**) Kyoto Encyclopedia of Genes and Genomes (KEGG) pathway analysis of DEGs. (**B**) Upregulated genes KEGG pathway analysis. (**C**) Downregulated genes KEGG pathway analysis.

**Figure S9 (related to Figure 4).**
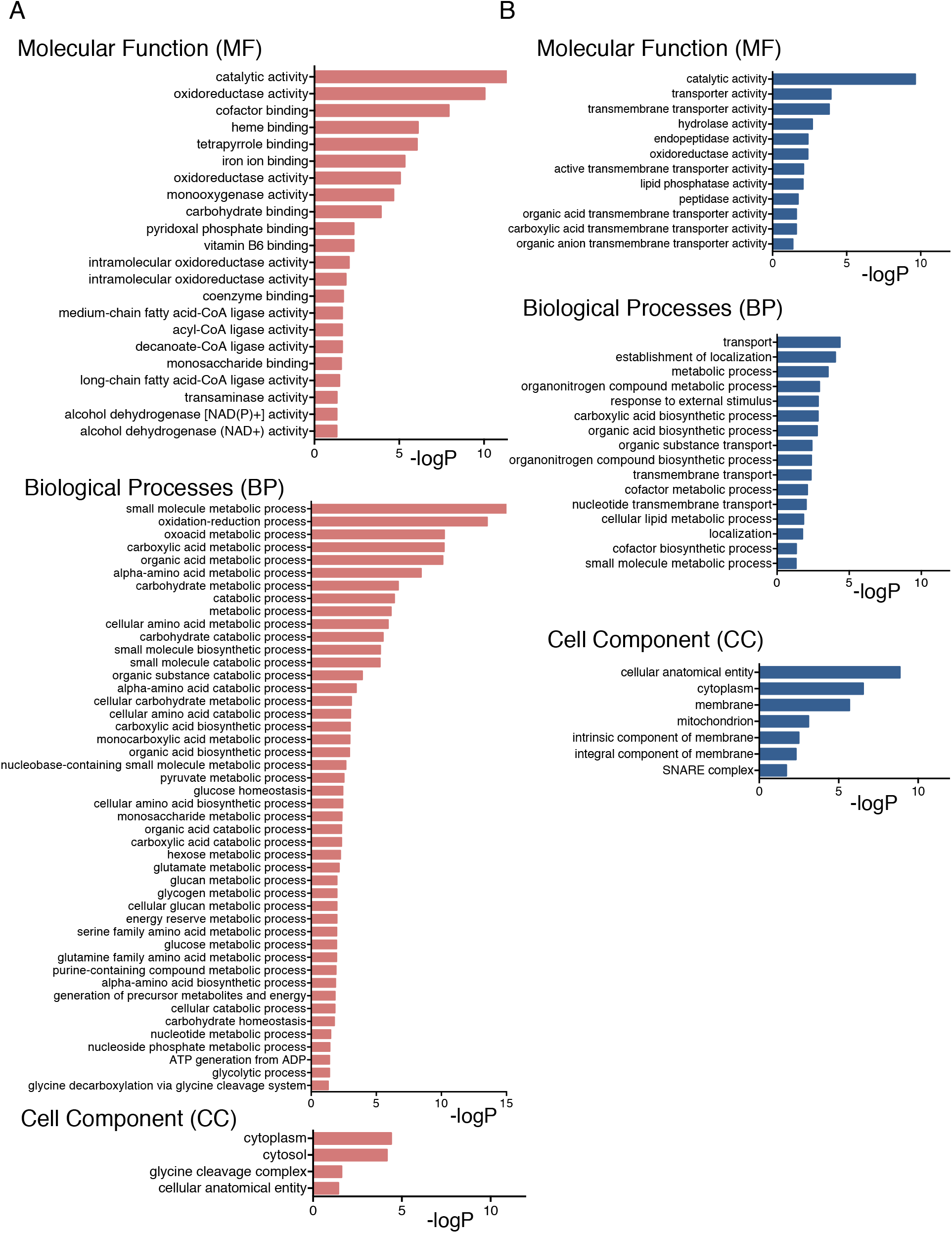
Enriched gene ontology (GO) functions of differentially expressed genes (DEGs) from fat body TFAM knock down mRNA Seq analysis. (**A**) Upregulated genes GO analysis. (**B**) Downregulated genes GO analysis.

**Figure S10 (related to Figure 4).**
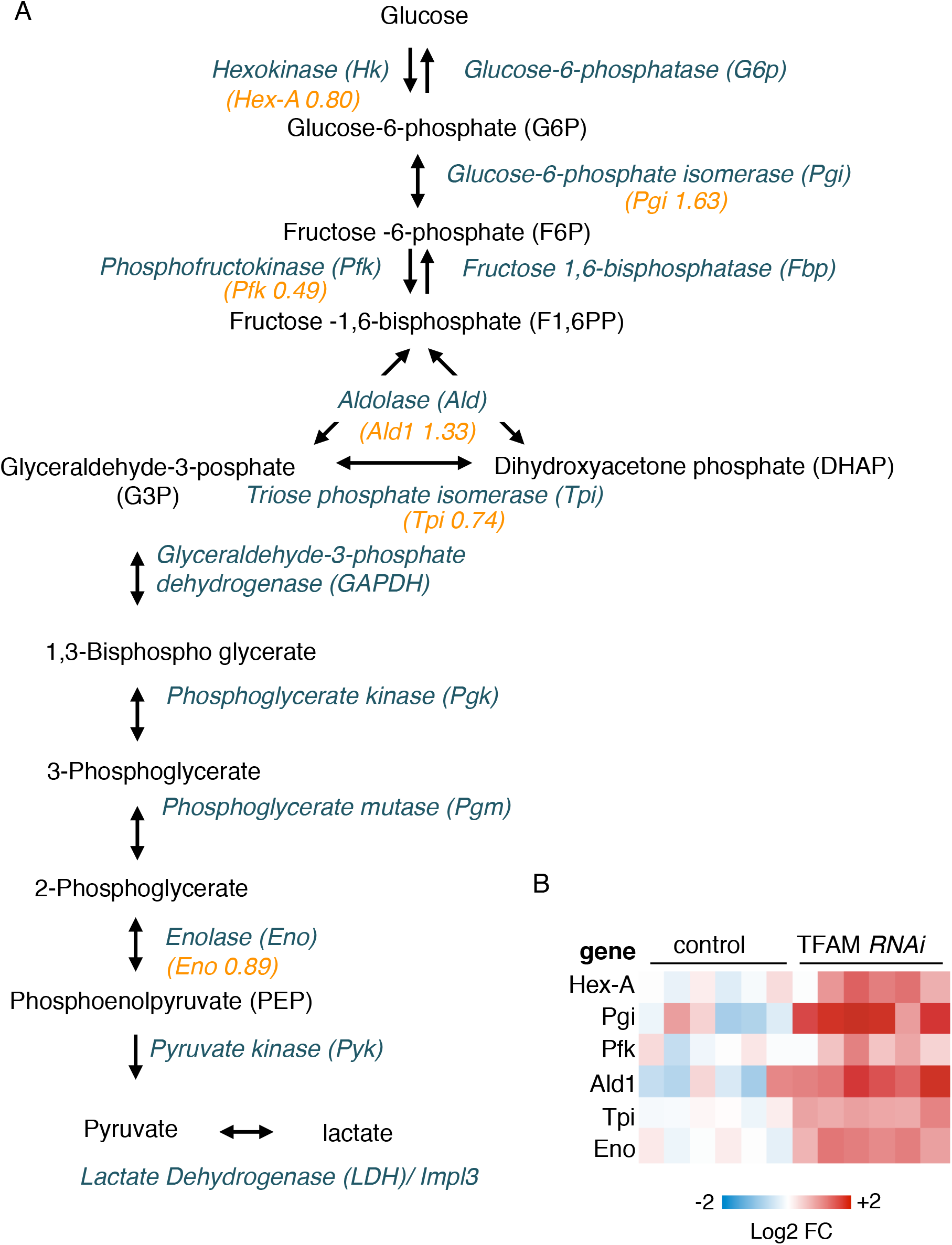
Glucose metabolism genes are upregulated in fat body specific TFAM knock down (RNAi #1). (**A**) A schematic of glycolytic pathway with relative fold change of mRNA indicated in orange color. (**B**) Heat maps of mRNA levels of the indicated mitochondrial genes in fat body isolated from control and 96 hrs AEL larvae. Each box represents a different replicate of a group of 20 animals. Color indicates normalized log2 fold change based on mRNA Seq transcripts per million (TPM) data.

**Figure S11 (related Figure 4).**
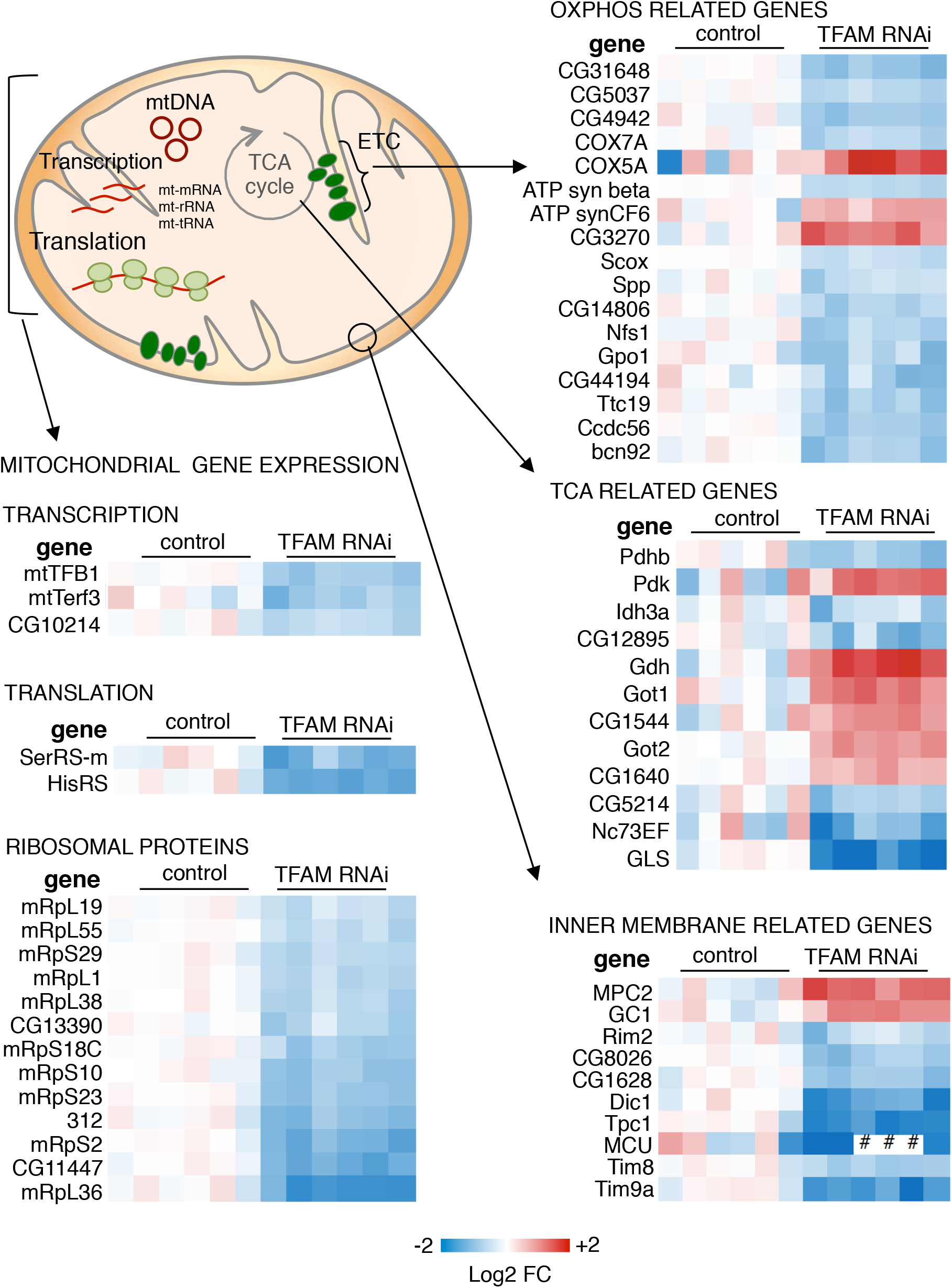
Mitochondrial genes in control and TFAM knock down (RNAi #1) fat body. Heat maps of mRNA levels of the indicated mitochondrial genes in fat body isolated from control and 96 hrs AEL larvae. Each box represents a different replicate of a group of 20 animals. Color indicates normalized log2 fold change based on mRNA Seq transcripts per million (TPM) data.

**Figure S12 (related to Figure 4).**
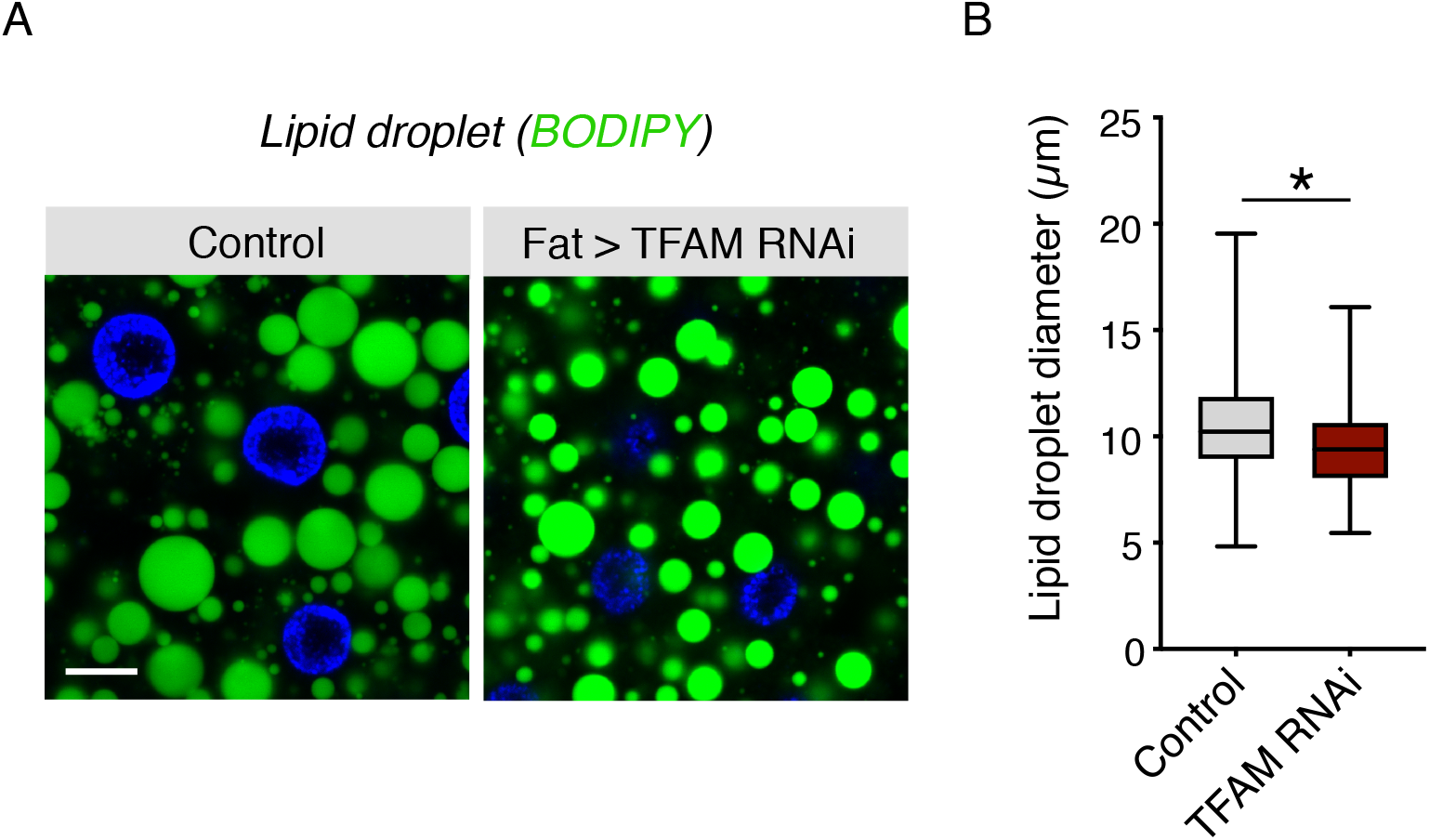
Lipid droplet size analysis in fat body with TFAM knock down (RNAi #1). Fat bodies were dissected from control or larvae expressing TFAM *RNAi* at 96 hrs AEL and stained with BODIPY to measure lipid droplet size. (**A-B**) Representative images and quantification of lipid droplet size. Data represented as mean ± SEM (* p < 0.05 unpaired t-test, ns = not significant.) Scale bar represents 10 μm.

**Figure S13 (related to Figure 4).**
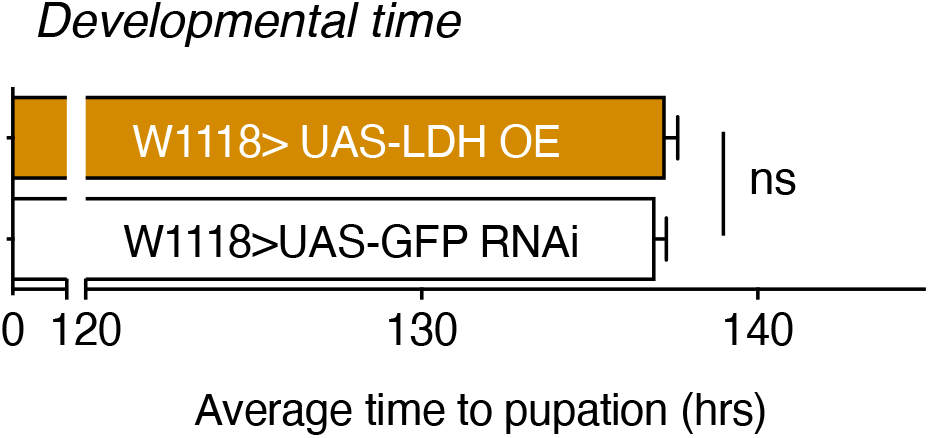
No driver control for LDH OE shows no change in the average time to pupation. n > 348 animals. Data represented as mean ± SEM (* p < 0.05 unpaired t-test, ns = not significant.)

**Supplementary Table 1:**
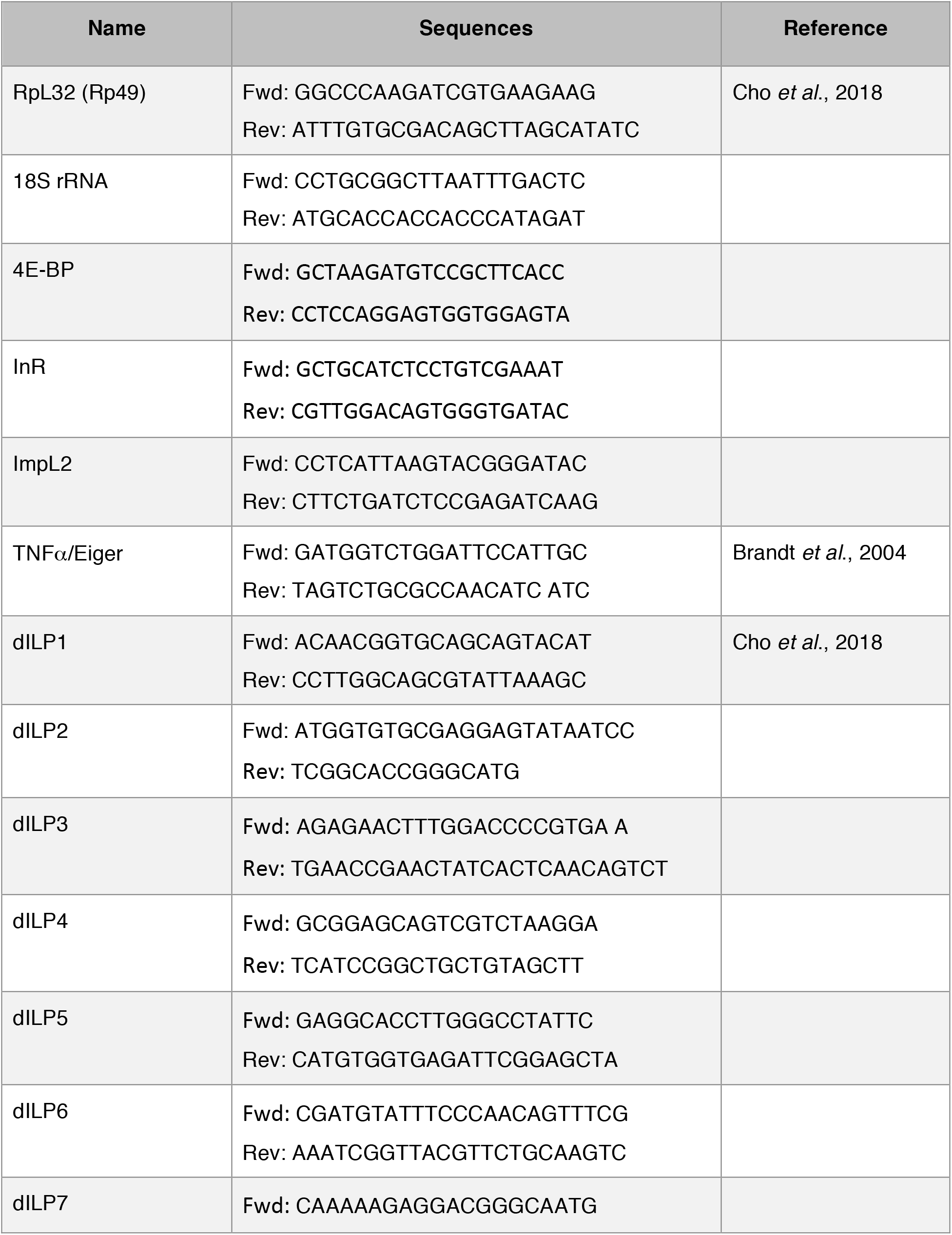
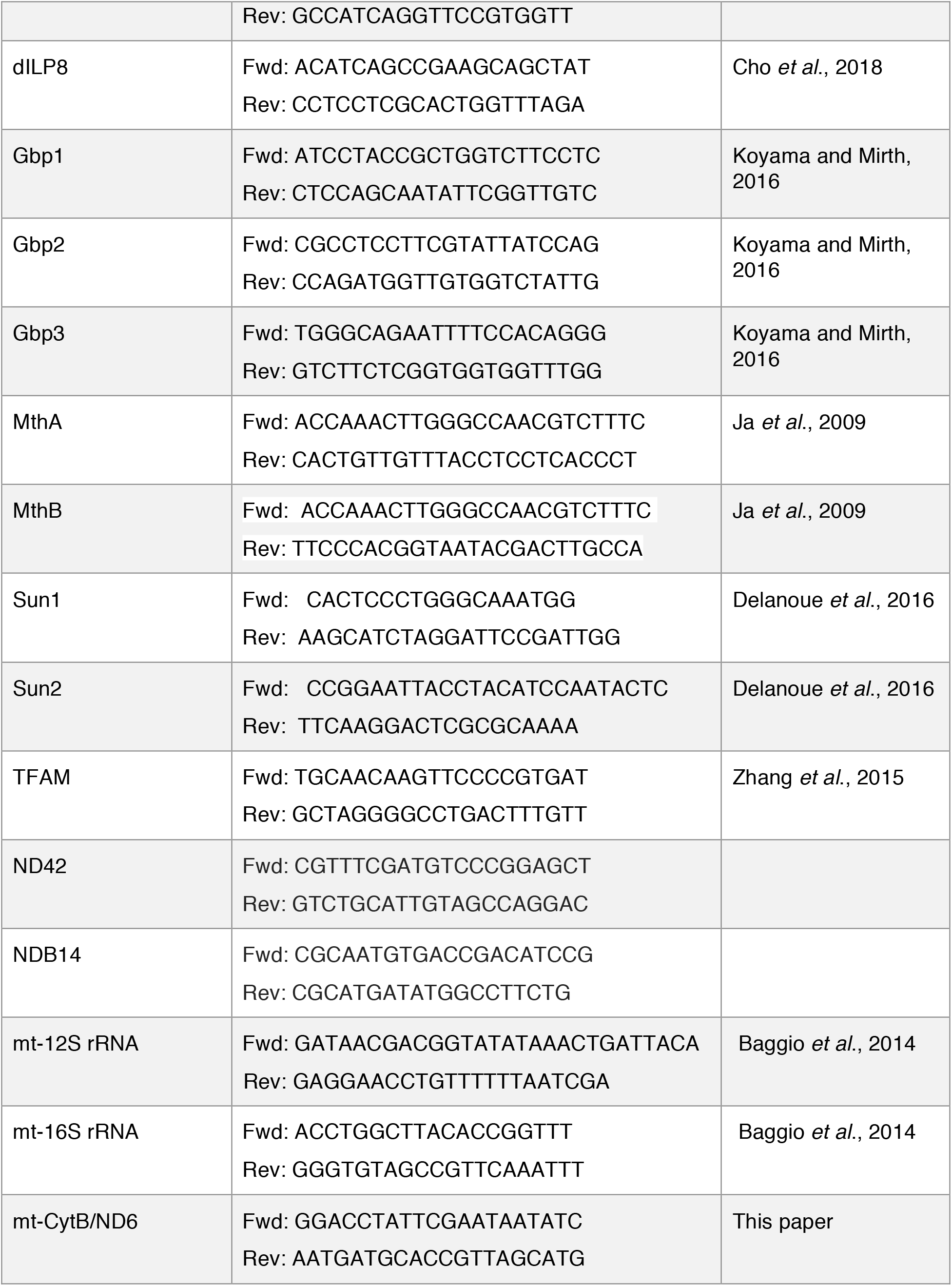
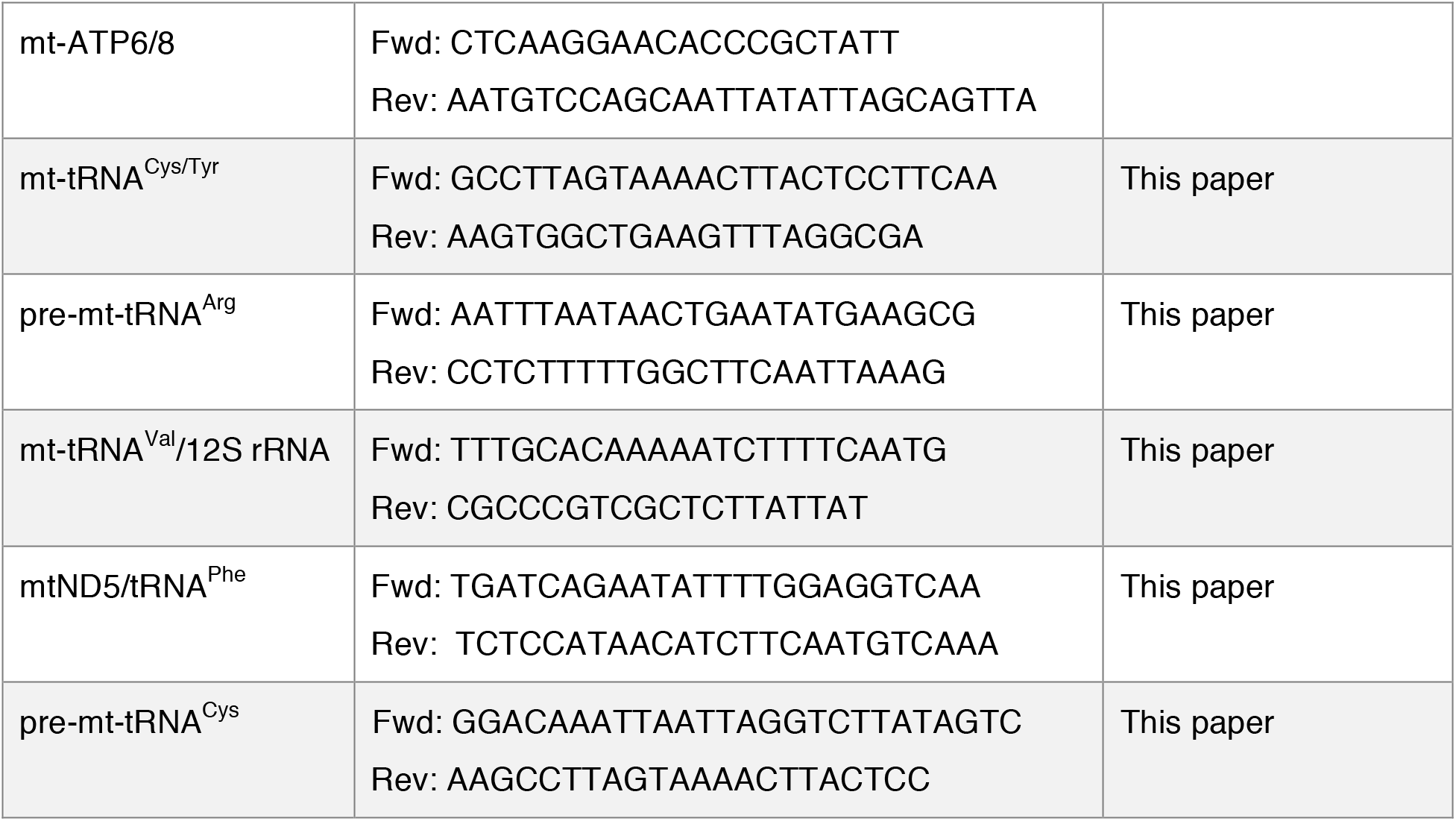
List of primer sequences for SYBR Green qRT-PCR

## References

Agrawal, N., Delanoue, R., Mauri, A., Basco, D., Pasco, M., Thorens, B., and Leopold, P. (2016). The Drosophila TNF Eiger Is an Adipokine that Acts on Insulin-Producing Cells to Mediate Nutrient Response. Cell metabolism 23, 675–684.

Andersen, D.S., Colombani, J., and Leopold, P. (2013). Coordination of organ growth: principles and outstanding questions from the world of insects. Trends in cell biology 23, 336–344.

Baksh, S.C., and Finley, L.W.S. (2021). Metabolic Coordination of Cell Fate by alpha-Ketoglutarate-Dependent Dioxygenases. Trends in cell biology 31, 24–36.

Boulan, L., Milan, M., and Leopold, P. (2015). The Systemic Control of Growth. Cold Spring Harb Perspect Biol 7.

Bricker, D.K., Taylor, E.B., Schell, J.C., Orsak, T., Boutron, A., Chen, Y.C., Cox, J.E., Cardon, C.M., Van Vranken, J.G., Dephoure, N., et al. (2012). A mitochondrial pyruvate carrier required for pyruvate uptake in yeast, Drosophila, and humans. Science (New York, NY) 337, 96–100.

Chen, Q., Kirk, K., Shurubor, Y.I., Zhao, D., Arreguin, A.J., Shahi, I., Valsecchi, F., Primiano, G., Calder, E.L., Carelli, V., et al. (2018). Rewiring of Glutamine Metabolism Is a Bioenergetic Adaptation of Human Cells with Mitochondrial DNA Mutations. Cell metabolism 27, 1007–1025 e1005.

Chowdhary, S., Madan, S., Tomer, D., Mavrakis, M., and Rikhy, R. (2020). Mitochondrial morphology and activity regulate furrow ingression and contractile ring dynamics in Drosophila cellularization. Mol Biol Cell 31, 2331–2347.

DeBerardinis, R.J., and Chandel, N.S. (2016). Fundamentals of cancer metabolism. Sci Adv 2, e1600200.

DeBerardinis, R.J., and Chandel, N.S. (2020). We need to talk about the Warburg effect. Nat Metab 2, 127–129.

Delanoue, R., Meschi, E., Agrawal, N., Mauri, A., Tsatskis, Y., McNeill, H., and Leopold, P. (2016). Drosophila insulin release is triggered by adipose Stunted ligand to brain Methuselah receptor. Science (New York, NY) 353, 1553–1556.

Delanoue, R., Slaidina, M., and Leopold, P. (2010). The steroid hormone ecdysone controls systemic growth by repressing dMyc function in Drosophila fat cells. Developmental cell 18, 1012–1021.

Droujinine, I.A., and Perrimon, N. (2016). Interorgan Communication Pathways in Physiology: Focus on Drosophila. Annual review of genetics 50, 539–570.

Drummond-Barbosa, D., and Tennessen, J.M. (2020). Reclaiming Warburg: using developmental biology to gain insight into human metabolic diseases. Development 147.

Gandara, L., and Wappner, P. (2018). Metabo-Devo: A metabolic perspective of development. Mech Dev 154, 12–23.

Gillette, C.M., Tennessen, J.M., and Reis, T. (2021). Balancing energy expenditure and storage with growth and biosynthesis during Drosophila development. Developmental biology.

Grewal, S.S. (2009). Insulin/TOR signaling in growth and homeostasis: a view from the fly world. Int J Biochem Cell Biol 41, 1006–1010.

Grewal, S.S. (2012). Controlling animal growth and body size - does fruit fly physiology point the way? F1000 Biol Rep 4, 12.

Hackenbrock, C.R. (1968). Ultrastructural bases for metabolically linked mechanical activity in mitochondria. II. Electron transport-linked ultrastructural transformations in mitochondria. J Cell Biol 37, 345–369.

Hamanaka, R.B., Weinberg, S.E., Reczek, C.R., and Chandel, N.S. (2016). The Mitochondrial Respiratory Chain Is Required for Organismal Adaptation to Hypoxia. Cell reports 15, 451–459.

Homem, C.C., Repic, M., and Knoblich, J.A. (2015). Proliferation control in neural stem and progenitor cells. Nat Rev Neurosci 16, 647–659.

Honegger, B., Galic, M., Kohler, K., Wittwer, F., Brogiolo, W., Hafen, E., and Stocker, H. (2008). Imp-L2, a putative homolog of vertebrate IGF-binding protein 7, counteracts insulin signaling in Drosophila and is essential for starvation resistance. J Biol 7, 10.

Hotamisligil, G.S., Arner, P., Caro, J.F., Atkinson, R.L., and Spiegelman, B.M. (1995). Increased adipose tissue expression of tumor necrosis factor-alpha in human obesity and insulin resistance. J Clin Invest 95, 2409–2415.

Hotamisligil, G.S., Shargill, N.S., and Spiegelman, B.M. (1993). Adipose expression of tumor necrosis factor-alpha: direct role in obesity-linked insulin resistance. Science (New York, NY) 259, 87–91.

Hotamisligil, G.S., and Spiegelman, B.M. (1994). Tumor necrosis factor alpha: a key component of the obesity-diabetes link. Diabetes 43, 1271–1278.

Hu, B., Li, H., and Zhang, X. (2021). A Balanced Act: The Effects of GH-GHR-IGF1 Axis on Mitochondrial Function. Front Cell Dev Biol 9, 630248.

Igaki, T., and Miura, M. (2014). The Drosophila TNF ortholog Eiger: emerging physiological roles and evolution of the TNF system. Semin Immunol 26, 267–274.

Ingaramo, M.C., Sanchez, J.A., Perrimon, N., and Dekanty, A. (2020). Fat Body p53 Regulates Systemic Insulin Signaling and Autophagy under Nutrient Stress via Drosophila Upd2 Repression. Cell reports 33, 108321.

John, G.B., Shang, Y., Li, L., Renken, C., Mannella, C.A., Selker, J.M., Rangell, L., Bennett, M.J., and Zha, J. (2005). The mitochondrial inner membrane protein mitofilin controls cristae morphology. Mol Biol Cell 16, 1543–1554.

Kanda, H., Igaki, T., Okano, H., and Miura, M. (2011). Conserved metabolic energy production pathways govern Eiger/TNF-induced nonapoptotic cell death. Proc Natl Acad Sci U S A 108, 18977–18982.

Kang, D., Kim, S.H., and Hamasaki, N. (2007). Mitochondrial transcription factor A (TFAM): roles in maintenance of mtDNA and cellular functions. Mitochondrion 7, 39–44.

Keller, L.C., Cheng, L., Locke, C.J., Muller, M., Fetter, R.D., and Davis, G.W. (2011). Glial-derived prodegenerative signaling in the Drosophila neuromuscular system. Neuron 72, 760–775.

Kelley, D.E., Goodpaster, B.H., and Storlien, L. (2002). Muscle triglyceride and insulin resistance. Annu Rev Nutr 22, 325–346.

Khacho, M., Clark, A., Svoboda, D.S., Azzi, J., MacLaurin, J.G., Meghaizel, C., Sesaki, H., Lagace, D.C., Germain, M., Harper, M.E., et al. (2016). Mitochondrial Dynamics Impacts Stem Cell Identity and Fate Decisions by Regulating a Nuclear Transcriptional Program. Cell Stem Cell 19, 232–247.

Koliaki, C., and Roden, M. (2014). Do mitochondria care about insulin resistance? Mol Metab 3, 351–353.

Koliaki, C., and Roden, M. (2016). Alterations of Mitochondrial Function and Insulin Sensitivity in Human Obesity and Diabetes Mellitus. Annu Rev Nutr 36, 337–367.

Koyama, T., and Mirth, C.K. (2016). Growth-Blocking Peptides As Nutrition-Sensitive Signals for Insulin Secretion and Body Size Regulation. PLoS Biol 14, e1002392.

Koyama, T., Texada, M.J., Halberg, K.A., and Rewitz, K. (2020). Metabolism and growth adaptation to environmental conditions in Drosophila. Cell Mol Life Sci 77, 4523–4551.

Li, H., Chawla, G., Hurlburt, A.J., Sterrett, M.C., Zaslaver, O., Cox, J., Karty, J.A., Rosebrock, A.P., Caudy, A.A., and Tennessen, J.M. (2017). Drosophila larvae synthesize the putative oncometabolite L-2-hydroxyglutarate during normal developmental growth. Proc Natl Acad Sci U S A 114, 1353–1358.

Li, H., Rai, M., Buddika, K., Sterrett, M.C., Luhur, A., Mahmoudzadeh, N.H., Julick, C.R., Pletcher, R.C., Chawla, G., Gosney, C.J., et al. (2019). Lactate dehydrogenase and glycerol-3-phosphate dehydrogenase cooperatively regulate growth and carbohydrate metabolism during Drosophila melanogaster larval development. Development 146.

Mabery, E.M., and Schneider, D.S. (2010). The Drosophila TNF ortholog eiger is required in the fat body for a robust immune response. J Innate Immun 2, 371–378.

Marshall, L., Rideout, E.J., and Grewal, S.S. (2012). Nutrient/TOR-dependent regulation of RNA polymerase III controls tissue and organismal growth in Drosophila. EMBO J 31, 1916–1930.

Martinez-Reyes, I., and Chandel, N.S. (2020). Mitochondrial TCA cycle metabolites control physiology and disease. Nat Commun 11, 102.

Matoo, O.B., Julick, C.R., and Montooth, K.L. (2019). Genetic Variation for Ontogenetic Shifts in Metabolism Underlies Physiological Homeostasis in Drosophila. Genetics 212, 537–552.

Mills, E.L., Kelly, B., Logan, A., Costa, A.S.H., Varma, M., Bryant, C.E., Tourlomousis, P., Dabritz, J.H.M., Gottlieb, E., Latorre, I., et al. (2016). Succinate Dehydrogenase Supports Metabolic Repurposing of Mitochondria to Drive Inflammatory Macrophages. Cell 167, 457–470 e413.

Mirth, C.K., Saunders, T.E., and Amourda, C. (2021). Growing Up in a Changing World: Environmental Regulation of Development in Insects. Annu Rev Entomol 66, 81–99.

Mitra, K., Rikhy, R., Lilly, M., and Lippincott-Schwartz, J. (2012). DRP1-dependent mitochondrial fission initiates follicle cell differentiation during Drosophila oogenesis. J Cell Biol 197, 487–497.

Miyazawa, H., and Aulehla, A. (2018). Revisiting the role of metabolism during development. Development 145.

Morita, M., Gravel, S.P., Chenard, V., Sikstrom, K., Zheng, L., Alain, T., Gandin, V., Avizonis, D., Arguello, M., Zakaria, C., et al. (2013). mTORC1 controls mitochondrial activity and biogenesis through 4E-BP-dependent translational regulation. Cell metabolism 18, 698–711.

Noguchi, T., Koizumi, M., and Hayashi, S. (2011). Sustained elongation of sperm tail promoted by local remodeling of giant mitochondria in Drosophila. Current biology: CB 21, 805–814.

Owusu-Ansah, E., Song, W., and Perrimon, N. (2013). Muscle mitohormesis promotes longevity via systemic repression of insulin signaling. Cell 155, 699–712.

Parisi, F., Stefanatos, R.K., Strathdee, K., Yu, Y., and Vidal, M. (2014). Transformed epithelia trigger non-tissue-autonomous tumor suppressor response by adipocytes via activation of Toll and Eiger/TNF signaling. Cell reports 6, 855–867.

Rajan, A., Housden, B.E., Wirtz-Peitz, F., Holderbaum, L., and Perrimon, N. (2017). A Mechanism Coupling Systemic Energy Sensing to Adipokine Secretion. Developmental cell 43, 83–98 e86.

Rajan, A., and Perrimon, N. (2012). Drosophila cytokine unpaired 2 regulates physiological homeostasis by remotely controlling insulin secretion. Cell 151, 123–137.

Rideout, E.J., Marshall, L., and Grewal, S.S. (2012). Drosophila RNA polymerase III repressor Maf1 controls body size and developmental timing by modulating tRNAiMet synthesis and systemic insulin signaling. Proc Natl Acad Sci U S A 109, 1139–1144.

Ryan, D.G., Murphy, M.P., Frezza, C., Prag, H.A., Chouchani, E.T., O’Neill, L.A., and Mills, E.L. (2019). Coupling Krebs cycle metabolites to signalling in immunity and cancer. Nat Metab 1, 16–33.

Ryan, D.G., and O’Neill, L.A.J. (2020). Krebs Cycle Reborn in Macrophage Immunometabolism. Annu Rev Immunol 38, 289–313.

Sanchez, J.A., Mesquita, D., Ingaramo, M.C., Ariel, F., Milan, M., and Dekanty, A. (2019). Eiger/TNFalpha-mediated Dilp8 and ROS production coordinate intra-organ growth in Drosophila. PLoS genetics 15, e1008133.

Sano, H., Nakamura, A., Texada, M.J., Truman, J.W., Ishimoto, H., Kamikouchi, A., Nibu, Y., Kume, K., Ida, T., and Kojima, M. (2015). The Nutrient-Responsive Hormone CCHamide-2 Controls Growth by Regulating Insulin-like Peptides in the Brain of Drosophila melanogaster. PLoS genetics 11, e1005209.

Sarraf-Zadeh, L., Christen, S., Sauer, U., Cognigni, P., Miguel-Aliaga, I., Stocker, H., Kohler, K., and Hafen, E. (2013). Local requirement of the Drosophila insulin binding protein imp-L2 in coordinating developmental progression with nutritional conditions. Developmental biology 381, 97–106.

Schell, J.C., Wisidagama, D.R., Bensard, C., Zhao, H., Wei, P., Tanner, J., Flores, A., Mohlman, J., Sorensen, L.K., Earl, C.S., et al. (2017). Control of intestinal stem cell function and proliferation by mitochondrial pyruvate metabolism. Nat Cell Biol 19, 1027–1036.

Senos Demarco, R., Uyemura, B.S., D’Alterio, C., and Jones, D.L. (2019). Mitochondrial fusion regulates lipid homeostasis and stem cell maintenance in the Drosophila testis. Nat Cell Biol 21, 710–720.

Shulman, G.I. (2004). Unraveling the cellular mechanism of insulin resistance in humans: new insights from magnetic resonance spectroscopy. Physiology (Bethesda) 19, 183–190.

Sieber, M.H., and Spradling, A.C. (2017). The role of metabolic states in development and disease. Curr Opin Genet Dev 45, 58–68.

Sieber, M.H., Thomsen, M.B., and Spradling, A.C. (2016). Electron Transport Chain Remodeling by GSK3 during Oogenesis Connects Nutrient State to Reproduction. Cell 164, 420–432.

Szendroedi, J., Phielix, E., and Roden, M. (2011). The role of mitochondria in insulin resistance and type 2 diabetes mellitus. Nat Rev Endocrinol 8, 92–103.

Tannahill, G.M., Curtis, A.M., Adamik, J., Palsson-McDermott, E.M., McGettrick, A.F., Goel, G., Frezza, C., Bernard, N.J., Kelly, B., Foley, N.H., et al. (2013). Succinate is an inflammatory signal that induces IL-1beta through HIF-1alpha. Nature 496, 238–242.

Tennessen, J.M., Baker, K.D., Lam, G., Evans, J., and Thummel, C.S. (2011). The Drosophila estrogen-related receptor directs a metabolic switch that supports developmental growth. Cell metabolism 13, 139–148.

Tennessen, J.M., Bertagnolli, N.M., Evans, J., Sieber, M.H., Cox, J., and Thummel, C.S. (2014). Coordinated metabolic transitions during Drosophila embryogenesis and the onset of aerobic glycolysis. G3 (Bethesda) 4, 839–850.

Texada, M.J., Koyama, T., and Rewitz, K. (2020). Regulation of Body Size and Growth Control. Genetics 216, 269–313.

Vernochet, C., Mourier, A., Bezy, O., Macotela, Y., Boucher, J., Rardin, M.J., An, D., Lee, K.Y., Ilkayeva, O.R., Zingaretti, C.M., et al. (2012). Adipose-specific deletion of TFAM increases mitochondrial oxidation and protects mice against obesity and insulin resistance. Cell metabolism 16, 765–776.

Wredenberg, A., Wibom, R., Wilhelmsson, H., Graff, C., Wiener, H.H., Burden, S.J., Oldfors, A., Westerblad, H., and Larsson, N.G. (2002). Increased mitochondrial mass in mitochondrial myopathy mice. Proc Natl Acad Sci U S A 99, 15066–15071.

Zhang, G.F., Jensen, M.V., Gray, S.M., El, K., Wang, Y., Lu, D., Becker, T.C., Campbell, J.E., and Newgard, C.B. (2021). Reductive TCA cycle metabolism fuels glutamine- and glucose-stimulated insulin secretion. Cell metabolism 33, 804–817 e805.

